# An open-source multiple-bioreactor system for replicable gas-fermentation experiments: Nitrate feed results in stochastic inhibition events, but improves ethanol production of *Clostridium ljungdahlii* with CO_2_ and H_2_

**DOI:** 10.1101/2019.12.15.877050

**Authors:** Christian-Marco Klask, Nicolai Kliem-Kuster, Bastian Molitor, Largus T. Angenent

**Affiliations:** Environmental Biotechnology Group, Center for Applied Geoscience, University of Tübingen, 72074 Tübingen, Germany; Max Planck Fellow, Max Planck Institute for Developmental Biology, 72076 Tübingen, Germany

**Keywords:** gas fermentation, *Clostridium ljungdahlii*, acetogenic bacteria, bioreactor system, pH-regulation, nitrate

## Abstract

The pH-value in fermentation broth has a large impact on the metabolic flux and growth behavior of acetogens. A decreasing pH level throughout time due to undissociated acetic acid accumulation is anticipated under uncontrolled pH conditions such as in bottle experiment. As a result, the impact of changes in the metabolism (*e.g*., due to a genetic modification) might remain unclear or even unrevealed. In contrast, pH-controlled conditions can be easily achieved in commercially available bioreactors. However, their acquisition is costly and their operation is time consuming, and therefore the experiment is often limited to a single bioreactor run. Here, we present a self-built, relatively cheap, and easy to handle open-source multiple-bioreactor system (MBS) consisting of six pH-controlled bioreactors at a 1-L scale. The functionality of the MBS was tested in three experiments by cultivating the acetogen *Clostridium ljungdahlii* with CO_2_ and H_2_ at steady-state conditions (=chemostat). The experiments were addressing the questions: (1) does the MBS provide replicable data for gas-fermentation experiments?; (2) does feeding acetate alter the production rate of ethanol; and (3) does feeding nitrate influence the product spectrum under controlled pH conditions with CO_2_ and H_2_? We applied four different periods in each experiment ranging from pH 6.0 to pH 4.5. Our data show high reproducibility for gas-fermentation experiments with *C. ljungdahlii*, using the MBS. We found that feeding acetate did not improve ethanol production, but rather impaired growth and reduced acetate production. Using nitrate as sole N-source, on the other hand, enhanced biomass production even at a low pH. However, we observed differences in growth, acetate, and ethanol production rates between triplicate bioreactors (n=3). We explained the different performances because of stochastic inhibition events, which we observed through the accumulation of nitrite, and which led to complete crashes at different operating times. One of these bioreactors recovered after the crash and showed enhanced ethanol production rates while simultaneously producing less acetate. The MBS offers a great opportunity to perform bench-scale bioreactor experiments at steady-state conditions with replicates, which is especially attractive for academia.

## Introduction

An increasing world population will likely lead to growing energy demands. To meet these demands in a sustainable way, we need to rethink the *status quo* of a fossil-based economy and transition into a renewable-based and circular economy. Furthermore, we have to mitigate the apparent climate effects of anthropogenic greenhouse gas emissions, such as carbon dioxide (CO_2_), which are caused preliminary by industry, agriculture, and transportation. Biotechnology offers potential to contribute to climate-friendly and economically feasible solutions. One promising solution is synthesis gas (syngas) fermentation with microbes (Mohammadi et al., 2011). For syngas fermentation, mixtures of the gases CO_2_, hydrogen (H_2_), and carbon monoxide (CO) are converted into products, such as acetate and ethanol, by acetogenic bacteria (Dürre, 2017). This process provides a promising way to produce chemicals and biofuels with a reduced CO_2_-footprint (Latif et al., 2014; Molitor et al., 2017; Phillips et al., 2017).

In recent years, the company LanzaTech (Skokie, IL, USA) demonstrated that ethanol production from syngas with the acetogen *Clostridium autoethanogenum* is possible at commercial scale, which further indicates the potential of this platform. While the LanzaTech technology is based on proprietary strains of *C. autoethanogenum*, in academic research the most frequently studied acetogen is the closely related microbe *Clostridium ljungdahlii*. Both microbes produce acetic acid, ethanol, and some 2,3-butanediol from gaseous substrates (Tanner et al., 1993; Abrini et al., 1994; Köpke et al., 2010; Brown et al., 2014).

Different strategies are employed to optimize *C. autoethanogenum* and *C. ljungdahlii* for biotechnology. On the one hand, genetic engineering is used to generate modified strains that produce butyrate (Köpke et al., 2010), butanol (Köpke and Liew, 2012; Ueki et al., 2014), acetone, and isopropanol (Bengelsdorf et al., 2016; Köpke et al., 2016). In academic research, the physiological characterization of these genetically engineered strains is typically performed in batch experiments with serum bottles, which does not allow to control important process parameters such as the pH-value. On the other hand, bioprocess engineering is used to investigate and optimize the production of naturally occurring products, such as ethanol, in optimized bioreactor systems (Younesi et al., 2005; Mohammadi et al., 2012; Richter et al., 2013; Abubackar et al., 2015). While the impact of cultivation parameters can be investigated within one study, these studies often are difficult to compare with each other, because very different bioreactor architectures and process parameters are used (Asimakopoulos et al., 2018). Furthermore, because of the complexity, these setups are not suitable to perform preliminary experiments with genetically engineered strains. These issues can be partly overcome by utilizing commercially available bioreactor (chemostat) systems. However, these systems are costly, and therefore often not available to laboratories that do not focus on bioprocess engineering. Consequently, genetically engineered strains are typically not studied in fermentations beyond the serum bottle size, which leaves a gap between the construction of these strains and the investigation under controlled fermentation conditions.

To close this gap, we developed a cost-efficient, open-source multiple-bioreactor system (MBS) that can be built from off-the-shelf components at a considerably lower cost compared to the cost of commercial bioreactor systems. We give all information on purchasing the required parts, the assembly of the MBS, the process control elements (*e.g*., stirring, pH, temperature), and further improvement ideas. We tested our MBS with *C. ljungdahlii* and CO_2_ and H_2_ as substrate under controlled pH conditions by addressing three questions: (1) does the MBS provide replicable data for gas-fermentation experiments; (2) does feeding acetate alter the production rate of ethanol; and (3) does feeding nitrate influence the product spectrum?

We selected the first question to test the reproducibility of our system, while the latter two questions address the physiology of the microbe. Acetogens such as *C. ljungdahlii* are known to be highly dependent on the pH in the fermentation broth. Since their main fermentation product is acetate, which acts (in the form of the undissociated acetic acid) as a weak acid, no pH control in serum bottles would intrinsically lower the pH of the medium during growth, but can be controlled in bioreactors such as in our MBS (Drake et al., 2008).

Our second question addresses acetate as a fermentation product directly. It has been demonstrated that feeding acetate from the first stage to the second stage in a two-stage bioreactor system with syngas including CO led to increased ethanol production rates with *C. ljungdahlii* (Richter et al., 2013). Others have shown that feeding acetate to a single bioreactor with CO_2_ and H_2_ also led to increased ethanol production rates with *C. autoethanogenum* (Mock et al., 2015). In both cases the pH was an important parameter because ethanol production is thermodynamically triggered at a low pH (Richter et al., 2016).

Our third question addresses the finding from a recent study in which nitrate was used as an alternative electron acceptor by *C. ljungdahlii*, while it also served as sole nitrogen source (N-source) in ammonium-free medium (Emerson et al., 2019). The co-utilization of CO_2_ and nitrate enhanced the autotrophic biomass formation with CO_2_ and H_2_ compared to standard cultivation conditions with ammonium as the sole N-source. Contrarily, ethanol production was strongly reduced under nitrate conditions with CO_2_ and H_2_. The authors discussed that nitrate reduction consumes electrons, which would be no longer available for the production of ethanol. At the same time, nitrate reduction led to an accumulation of ammonium, which resulted in a continuous increase of the pH from 6.0 to 8.0 during the batch cultivations in serum bottles (Emerson et al., 2019), which would intrinsically prevent ethanol production. These two questions on the physiology address growth conditions in which the medium pH has a considerable impact on the production of biomass and fermentation products, such as acetate and ethanol, and therefore are perfectly suited to demonstrate the functionality of our MBS.

## Materials and Methods

### Microbial strains and medium composition

Wild type *C. ljungdahlii* PETC (DSM 13528) was obtained from the DSMZ (Braunschweig, Germany) and used for all experiments. Pre-cultures were grown heterotrophically at 37°C (IN260 stand incubator, Memmert, Germany) in 100-mL serum bottles with 50 mL of standard PETC medium containing (per liter): 0.5 g yeast extract; 1.0 g NH_4_Cl; 0.1 g KCl; 0.2 g MgSO_4_·7 H_2_O; 0.8 g NaCl; 0.1 g KH_2_PO_4_; 0.02 g CaCl_2_·2 H_2_O; 4 mL resazurin-solution (0.025 vol%); 10 ml trace element solution (TE, 100x); 10 mL Wolfe’s vitamin solution (100x); 10 mL reducing agent (100x); and 20 mL of fructose/2-(*N*-morpholino)ethanesulfonic acid (MES) solution (50x). Vitamins, reducing agent, and fructose/MES solution were added after autoclaving under sterile conditions. TE was prepared as 100x stock solution containing (per liter): 2 g nitrilotriacetic acid (NTA); 1 g MnSO_4_·H_2_O; 0.8 g Fe(SO_4_)_2_(NH_4_Cl)_2_·6 H_2_O; 0.2 g CoCl_2_·6 H_2_O; 0.0002 g ZnSO_4_·7 H_2_O; 0.2 g CuCl_2_·2 H_2_O; 0.02 g NiCl_2_·6 H_2_O; 0.02 g Na_2_MoO_4_·2 H_2_O; 0.02 g Na_2_SeO_4_; and 0.02 g Na_2_WO_4_. The pH of the TE was adjusted to 6.0 after adding NTA. The solution was autoclaved and stored at 4°C. Wolfe’s vitamin solution was prepared aerobically containing (per liter): 2 mg biotin; 2 mg folic acid; 10 mg pyridoxine-hydrochloride; 5 mg thiamin-HCl; 5 mg riboflavin; 5 mg nicotinic acid; 5 mg calcium pantothenate; 5 mg *p*-aminobenzoic acid; 5 mg lipoic acid; and 0.1 mg cobalamin. The vitamin solution was sterilized using a sterile filter (0.2 µm), sparged with N_2_ through a sterile filter, and stored at 4°C. The 50x fructose/MES solution contained (per 100 mL): 25 g fructose; and 10 g MES. The pH was adjusted to 6.0 by adding KOH. The solution was sterilized, sparged with N_2_ through a sterile filter, and stored at room temperature. The reducing agent was prepared under 100% N_2_ in a glove box (UniLab Pro Eco, MBraun, Germany) and contained (per 100 mL): 0.9 g NaOH; 4 g cysteine-HCl; and 2.17 g/L Na_2_S (60 weight%). Anaerobic water was used for the preparation of the reducing agent. The reducing agent was autoclaved and stored at 4°C.

For all bioreactor experiments, the standard PETC medium for the initial batch phase was supplemented with 0.5 g L^−1^ yeast extract and autoclaved inside the bioreactor vessel with an open off-gas line to enable pressure balance. The autoclaved bioreactors were slowly cooled down at room temperature overnight with an attached sterile filter at the off-gas line. After transferring each bioreactor to the MBS frame, the medium was continuously sparged with a gas mixture of CO_2_ and H_2_ (20:80 vol%) through a sterile filter. After 1 h, vitamins and reducing agent were added through the sampling port. N_2_ gas was applied through a sterile filter to flush the sampling port after each addition of media components. Subsequently, each bioreactor was inoculated with 5 mL of an exponential heterotrophically grown PETC culture (OD_600_ 0.5-0.8). Feed bottles containing 4 L of PETC medium with additions, as described below, were prepared for continuous mode, autoclaved, and stored overnight with an attached sterile filter on the off-gas line. The bottles were sparged with N_2_ for 2 h through a sterile filter. Vitamins and reducing agents were added under sterile conditions. A gas bag with N_2_ gas was attached with a sterile filter to balance the pressure in the feed bottle during the bioreactor run. Standard PETC medium for continuous mode did not contain yeast extract and was adjusted to the respective pH of the period. For the first experiment we used standard PETC medium. For the second experiment, we added 100 mM Na-acetate or 100 mM NaCl (to give a similar ion strength). For the third experiment, we supplied no NH_4_Cl, but we supplied the equivalent amount of nitrogen as NaNO_3_ (18.7 mM).

### Bioreactor setup and standard operating conditions

Six 1-L self-built bioreactors (**Fig. 1, Results S1**) with a working volume of 0.5 L were used for the three experimental bioreactor runs **(Fig. 2, Fig. 3, Fig. S4)**. The cultivation temperature was 37°C and the agitation was set to 300 rpm. The gas flow rate was adjusted to 30 mL min^−1^ *prior* to inoculation. For continuous-mode operating conditions, the medium feed rate was measured to be 0.10 mL min^−1^, which resulted in a 3.5-day hydraulic retention time (HRT), and which represents 1.7 HRT periods within each period of six days. To establish microbial growth in the MBS after one inoculation event for each bioreactor, we operated the MBS in batch mode for three days before switching to continuous mode. The pH was set to 6.0 during the batch mode and the first 6 days (Period I) in continuous mode. Subsequently, the pH setting was lowered stepwise in 6 days to a pH of 5.5, 5.0, and 4.5 (Period II-IV). While the pH of the feed medium was adjusted to the anticipated pH of the period, we did not use the acid feed to actively adjust the pH within the bioreactor. Instead, we let the pH decrease to the set value by the microbial production of undissociated acetic acid to avoid a pH shock. This took approximately two days of each 6-day period (**Table 1**). For the first experimental bioreactor run (**Fig. 2**), only base pumps were connected to the bioreactors. For the second experimental bioreactor run (**Fig. 3**), acid and base pumps were connected to the bioreactors with nitrate feed and only base pumps to the bioreactors with ammonium feed.

**Table 1.** Average values for OD_600_ and acetate/ethanol production rates during the continuous fermentation of *C. ljungdahlii* with CO_2_ and H_2_ at four different pH conditions in standard PETC medium using the MBS.

| Operating conditions | OD<sub>600</sub><sup>1</sup> | Acetate production rate<br>[mmol-C L<sup>-1</sup> d<sup>-1</sup>]<sup>1</sup> | Ethanol production rate<br>[mmol-C L<sup>-1</sup> d<sup>-1</sup>]<sup>1</sup> | Ratio<sub>Et/Ac</sub><sup>2</sup> |
| --- | --- | --- | --- | --- |
| Period I (pH=6.0) | 0.82 ± 0.04 | 73.1 ± 2.1 | 6.4 ± 1.6 | 0.1 |
| Period II (pH=5.5) | 0.69 ± 0.01 | 60.1 ± 3.4 | 22.5 ± 0.5 | 0.4 |
| Period III (pH=5.0) | 0.51 ± 0.05 | 53.7 ± 3.3 | 18.7 ± 4.8 | 0.3 |
| Period IV (pH=4.5) | 0.18 ± 0.06 | 29.2 ± 10.0 | 3.4 ± 1.4 | 0.1 |

<sup>1</sup> Values for the bioreactors with ammonium feed (n=3) are given as the average (± standard deviation) from three bioreactors for the last 5 data points of every period.
<sup>2</sup> Et, Ethanol; Ac, Acetate.

**Figure 1.**
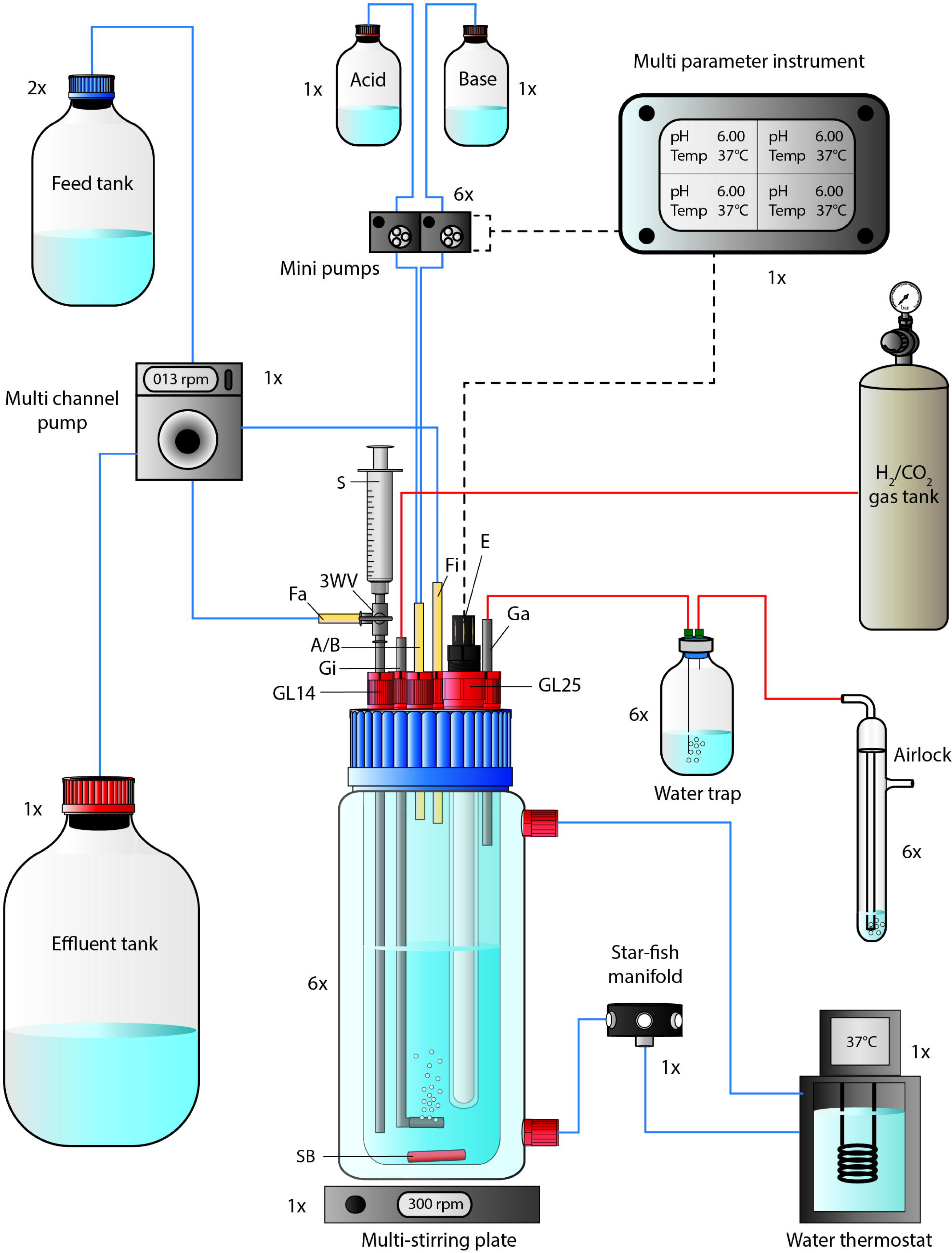
Flow chart of a single bioreactor operated in the MBS. The 1-L bioreactor vessel consisted of a double-walled glass vessel and a customized lid, while it was placed on a multi-stirring plate with up to six bioreactors. The bioreactor temperature was maintained through a water circulation unit at 37°C. The autoclavable lid offered connections for 5x GL14 and 1x GL25. A set of stainless-steel tubing was used for the gas-in/-out lines and for the medium feed-out line. The three-way valve at the medium feed-out line was required for sampling using a 5-mL syringe. The pH and bioreactor medium temperature was tracked *via* a pH/pt1000-electrode that was connected to a multi-parameter instrument. The multi-parameter instrument controlled and triggered two mini pumps (for base and acid) at programmable conditions. For continuous mode, the feed medium to each bioreactor was pumped *via* a single multi-channel pump from the feed tank into the bioreactor. The same pump was used to transfer the effluent from each bioreactor into the effluent tank. Sterile CO_2_ and H_2_ gas (20:80 vol-%) was sparged into the system through stainless-steel tubing with an attached sparger. The gas-out line was connected to a 100-mL serum bottle to serve as a water trap before the outgoing gas passed an airlock. The 1x, 2x, and 6x next to each unit in the figure describe the quantity, which is required to operate six bioreactors simultaneously. Abbreviations: A/B, Acid and/or base feed line; E, pH/pt1000 electrode; Fa, medium feed-out line; Fi, medium feed-in line; Ga, gas-out line; Gi, gas-in line; GL14, screw joint connection size 14; GL25, screw joint connection size 25; rpm, revolutions per minute; SB, stirring bar; 3WV, three-way valve. Blue lines indicate liquid transfer, red lines contain gas, and dotted black lines provide electric power or signals.

**Figure 2.**
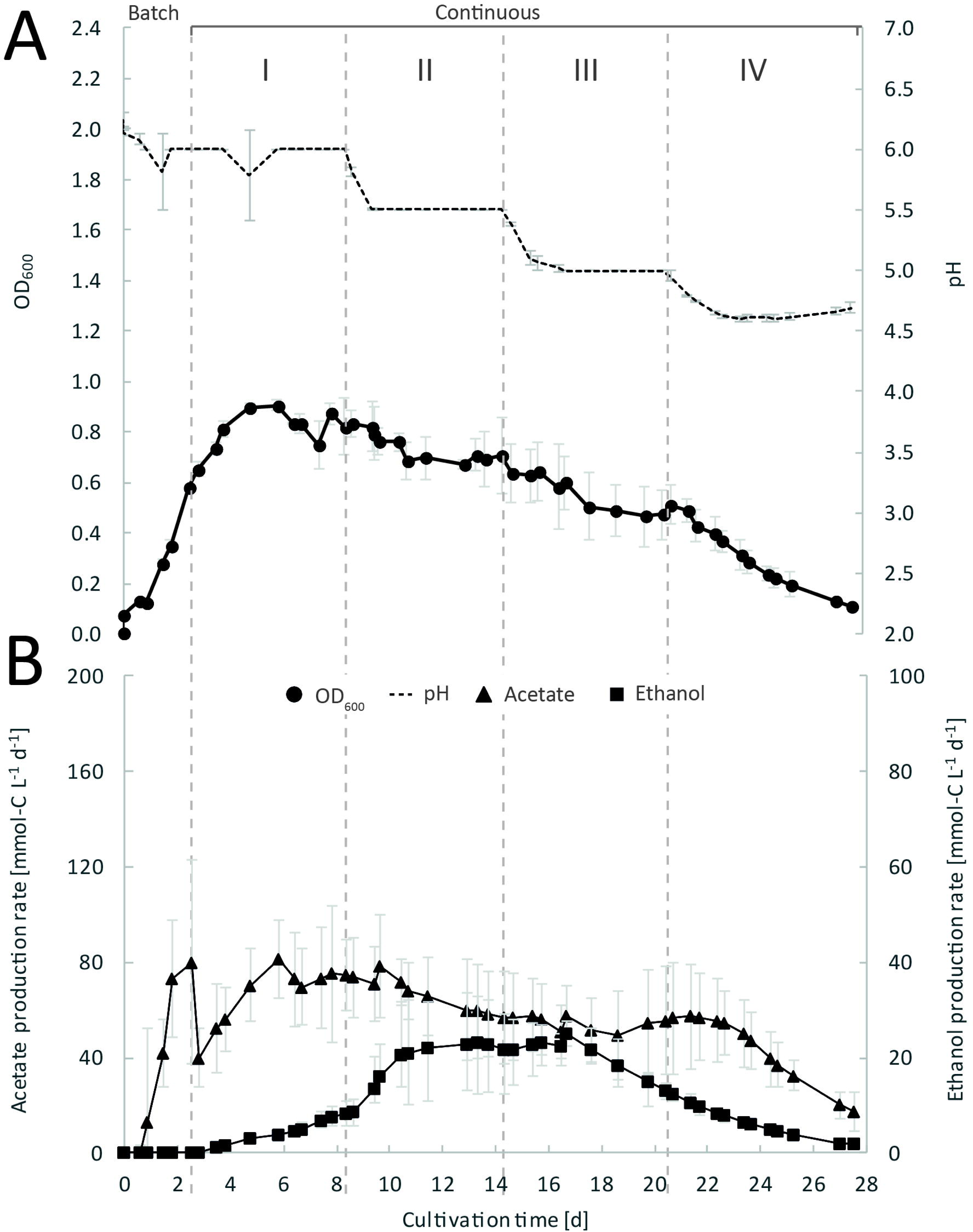
Continuous gas fermentation of *C. ljungdahlii* with CO_2_ and H_2_ in standard PETC medium at different periods. Mean values of triplicates with standard deviation (n=3) for pH and OD_600_ (A), and for acetate and ethanol production rates in mmol-C L^−1^ d^−1^ (B). Standard PETC medium containing 18.7 mM ammonium chloride as sole N-source was used. The horizontal dotted lines indicate the continuous process in which medium of different pH was fed to each bioreactor. Period: I, pH=6.0; II, pH=5.5, III, pH=5.0; and IV, pH=4.5.

**Figure 3.**
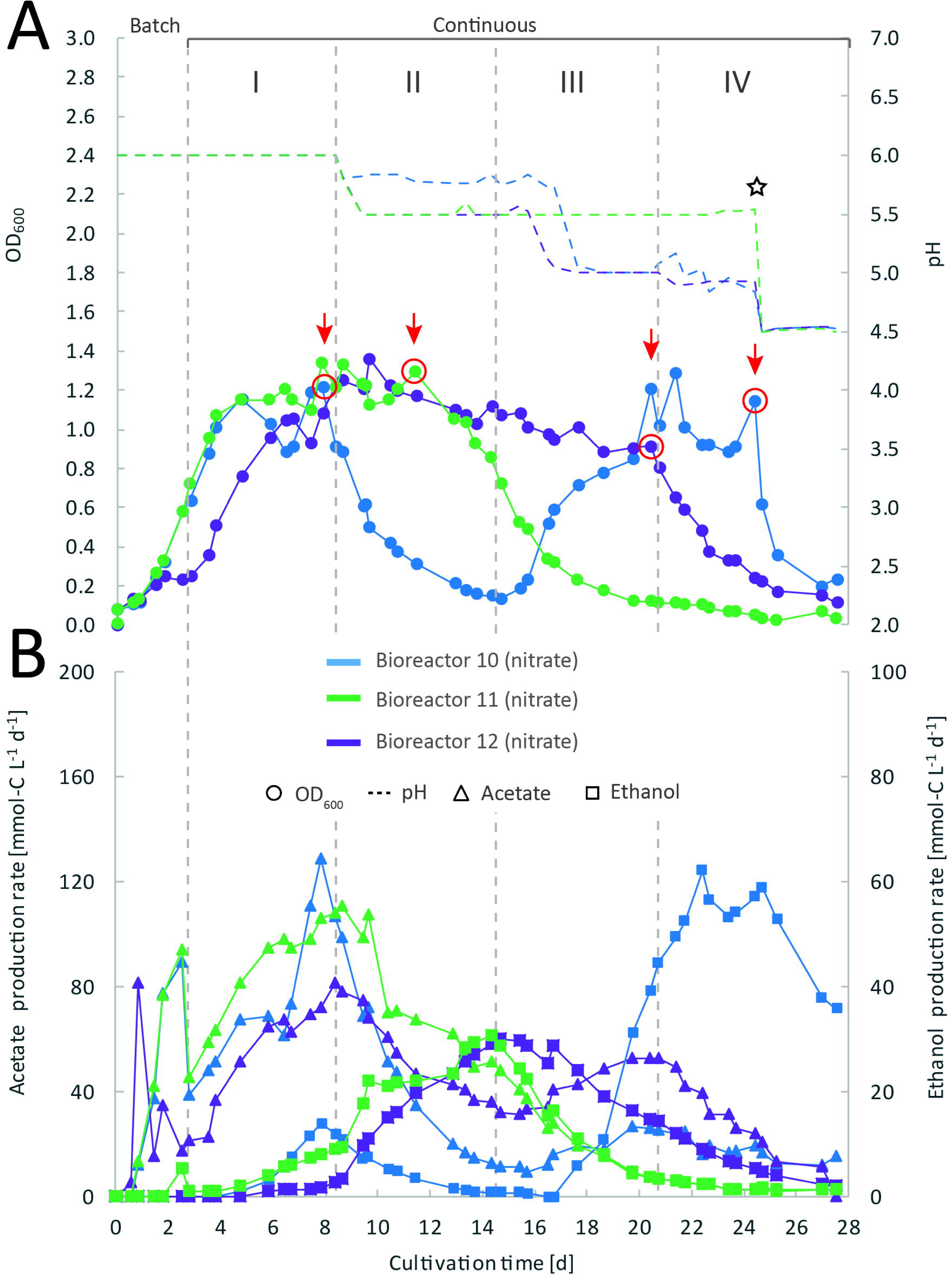
Impact of nitrate as alternative N-source on continuous gas fermentation of *C. ljungdahlii* using CO_2_ and H_2_ at different periods. Single values for pH and OD_600_ (A), and for acetate and ethanol production rates in mmol-C L^−1^ d^−1^ (B). The bioreactors with nitrate feed were grown in ammonium-free PETC medium supplemented with 18.7 mM Na-nitrate. The horizontal dotted lines indicate the continuous process in which medium of different pH was fed to each bioreactor. The red arrows and circles indicate the crash in OD_600_ of each bioreactor with nitrate feed at different time points. The star symbol describes the time point were the pH was lowered manually by adding HCl to the system until a pH of 4.5 was reached. Period: I, pH=6.0; II, pH=5.5, III, pH=5.0; and IV, pH=4.5.

### Sampling and analyses

Bioreactors were sampled once or twice per day. A pre-sample of 3 mL of cell suspension was discarded, before transferring 2 mL of cell suspension into 2-mL reaction tubes (main sample). The multi-channel pump was switched off during the sampling procedure. Cell growth was monitored by measuring the optical density at 600 nm (OD_600_) (Nanophotometer NP80, Implen, Germany). For OD_600_-values larger than 0.5, dilutions with 100 mM phosphate-buffered saline (PBS) at pH=7.4 were prepared. Nitrate and nitrite concentrations were qualitatively monitored using test stripes (Quantofix nitrate/nitrite, Macherey-Nagel, Germany). A correlation between cellular dry weight (CDW) and OD_600_ was calculated by harvesting 50 mL of culture sample from every bioreactor, centrifugation of the samples at 3428 relative centrifugal force (rcf) (Eppendorf centrifuge 5920R) for 12 min at room temperature (RT) and, subsequently, drying the pellet at 65°C for 3 days. The CDW for an OD_600_ of 1 was determined to be 0.24 g L^−1^ for cultures grown in PETC medium and 0.29 g L^−1^ for cultures grown in PETC medium with nitrate instead of ammonium.

Acetate and ethanol concentrations were analyzed *via* a high pressure liquid chromatography (HPLC) (LC20, Shimadzu, Japan) system that was equipped with an Aminex HPX-87H column and operated with 5 mM sulfuric acid as eluent. The flow was 0.6 mL min^−1^ (LC-20AD). The oven temperature was 65°C (CTO-20AC). The sample rack of the HPLC was constantly cooled to 15°C in the autosampler unit (SIL-20AC_HT_). For HPLC sample preparation, all culture samples were centrifuged for 3 min at 15871 rcf (Centrifuge 5424, Eppendorf, Germany) in 1.5-mL reaction tubes. 750 µl of the supernatant was transferred into clean reaction tubes and stored at -20°C until use. Frozen samples were thawed at 30°C and 250 revolutions per minute (rpm) for 10 min (Thermomixer C, Eppendorf, Germany). The samples were centrifuged again and 500 µl of the supernatant was transferred into short thread HPLC/GC vials (glass vial ND9, VWR, Germany) and sealed with short screw caps, which contained rubber septa (6 mm for ND9, VWR, Germany). New standards for acetate and ethanol were prepared for every analysis (retention time of acetate was 14.8 min and retention time of ethanol was 21.5 min). All samples were randomized.

## Results

### Designing and testing the functionality of the MBS

We based our experiments in this study on a versatile self-built multiple-bioreactor system (MBS). The MBS **(Fig. 1, Results S1, Fig. S1 and S2, Table S1)** was designed to either perform heterotrophic or autotrophic cultivation experiments in batch or continuous mode. The MBS can be used to operate up to six bioreactors simultaneously, each individually at different pH conditions or, if necessary, with different feed medium.

To show high comparability and reproducibility of our MBS, we grew *C. ljungdahlii* simultaneously as triplicates with standard PETC medium and CO_2_ and H_2_ (ammonium, bioreactors 1/2/3) during the first experiment (**Fig. 2, Fig. S3**). We observed that growth was similar in the triplicate bioreactors during the cultivation for 27.6 days. During the initial batch mode, the average OD_600_ increased to 0.58 ± 0.01 (**Fig. 2A**). After switching to continuous mode, the average OD_600_ increased further to values of 0.82 ± 0.04 during Period I. For Periods II, III, and IV, the average OD_600_ for the bioreactors constantly decreased to values of 0.69 ± 0.01, 0.51 ± 0.05, and 0.18 ± 0.06 (**Fig. 2A, Table 1**). As expected, the pH of each bioreactor was decreasing during all periods by microbial acetate production. The simultaneous and constant decrease of OD_600_ indicated reduced growth rates of *C. ljungdahlii* at a lower pH level in our system. In batch mode, the acetate production rates increased with increasing OD_600_, but then considerably dropped after switching to continuous mode (**Fig. 2A**). The acetate production rates increased again to an average value of 73.1 ± 2.1 mmol-C L^−1^ d^−1^ for Period I (**Fig. 2B**). The acetate production rates decreased to average values of 60.1 ± 3.4 mmol-C L^−1^ d^−1^, and 53.7 ± 3.3 mmol-C L^−1^ d^−1^ for Periods II and III, respectively. For Period IV, the acetate production rate had only an average value of 29.2 ± 10.0 mmol-C L^−1^ d^−1^ (**Fig. 2B**). Ethanol production rates in mmol-C L^−1^ d^−1^ were negligible during batch mode, but slowly increased after switching to continuous mode. The highest ethanol production rates were observed for Period II with average values of 22.5 ± 0.5 mmol-C L^−1^ d^−1^. During the Periods III and IV, the ethanol production rates kept decreasing to average values of 18.7 ± 4.8 mmol-C L^−1^ d^−1^ and 3.4 ± 1.4 mmol-C L^−1^ d^−1^, respectively (**Fig. 2B**). The results showed high reproducibility with small standard deviation for all tested parameters using the MBS.

### Feeding additional acetate to continuous and pH-controlled gas fermentation of *C. ljungdahlii* with CO_2_ and H_2_

During the second experiment, we investigated the impact of feeding acetate on the production of ethanol from CO_2_ and H_2_ during an operating period of 27.2 days **(Results S2, Fig. S4)**. For this, three bioreactors (Na-acetate, bioreactors 4/5/6) were fed with PETC medium that was supplemented with 100 mM Na-acetate, while three bioreactors (NaCl, bioreactors 7/8/9) were fed with PETC medium that was supplemented with 100 mM NaCl to achieve a similar ionic strength. We found again that our triplicate bioreactors were reproducible in terms of growth. However, feeding acetate did not result in the expected increased ethanol production rates. Instead, the additional acetate led to impaired growth, lower OD_600_-values, and reduced production rates of acetate and ethanol (**Fig. S4, Table S2**). However, feeding additional NaCl compared to our first experiment with standard PETC medium had a positive stimulating impact on ethanol production rates at lower pH (**Fig. S5 and S7, Table S2**). Growth (OD_600_) and acetate production were not influenced by the higher NaCl content in the medium and showed similarities to our preliminary experiment with standard PETC medium (**Fig. 2, Table 1**). The second experiment shows that acetate did not, but that NaCl did improve ethanol production of *C. ljungdahlii* with CO_2_ and H_2_.

### Applying nitrate as a sole N-source to a continuous and pH-controlled gas fermentation of *C. ljungdahlii* with CO_2_ and H_2_

During our third experiment with an operating period of 27.5 days, we investigated the impact of nitrate as an alternative N-source on the production of ethanol from CO_2_ and H_2_ **(Fig. 3)**. For this experiment, three bioreactors (nitrate, bioreactors 10/11/12) were fed with PETC medium containing nitrate instead of ammonium at an equivalent molar amount of nitrogen (=18.7 mM). We found an increasing pH due to ammonium production in preliminary bottle experiments in nitrate-containing PETC medium (**Fig. S7**). A pH increase was also observed in the nitrate bottle experiments of Emerson et al. (2019). Despite the pH-control in our experiment, all bioreactors with nitrate feed showed remarkable differences in growth, pH, acetate production, and ethanol production rates. Therefore, we report individual data for each bioreactor and highlight lowest and highest values (**Fig. 3, Table 2**). We use the data of the first experiment (ammonium, bioreactor 1/2/3) as the control in which ammonium served as the sole N-source (**Fig. 2, Table 1**). Unexpectedly, we observed a pH-buffering effect in all bioreactors with nitrate feed during the fermentation. This was most likely due to an interplay between the produced acetate and ammonium by the microbes. Overall, the pH was slowly decreasing in all bioreactors with nitrate feed (**Fig. 3A**), and we did not measure increasing pH values.

**Table 2.** Highest observed values for OD_600_ and acetate/ethanol production rates at specific pH during continuous fermentation of *C. ljungdahlii* with CO_2_/H_2_ and nitrate.

| Highest value for | Bioreactor 10<br>(nitrate) | Bioreactor 11<br>(nitrate) | Bioreactor 12<br>(nitrate) | Bioreactor 1-3<br>(ammonium)<sup>1</sup> |
| --- | --- | --- | --- | --- |
| OD<sub>600</sub> | 1.29<br>(pH 5.2) | 1.36<br>(pH 5.5) | 1.34<br>(pH 6.0) | 0.90 ± 0.02<br>(pH 6.0) |
| Acetate production rate<br>[mmol-C L<sup>-1</sup> d<sup>-1</sup>] | 128.8<br>(pH 6.0) | 81.5<br>(pH 6.0) | 108.3<br>(pH 6.0) | 81.4 ± 3.0<br>(pH 6.0) |
| Ethanol production rate<br>[mmol-C L<sup>-1</sup> d<sup>-1</sup>] | 62.0<br>(pH 5.0) | 29.9<br>(pH 5.0) | 30.6<br>(pH 5.5) | 25.0 ± 2.7<br>(pH 5.0) |
| Ratio<sub>Et/Ac</sub><sup>2</sup> | 3.8<br>(pH 4.9) | 0.7<br>(pH 5.5) | 0.6<br>(pH 5.5) | 0.4<br>(pH 5.5) |

<sup>1</sup> Values for the bioreactors with ammonium feed (n=3) are given as the average (± standard deviation) from three bioreactors for the last 5 data points.
<sup>2</sup> Et, Ethanol; Ac, Acetate.

During the initial batch mode, two of the bioreactors with nitrate feed (bioreactor 10 and 11) reached OD_600_-values of 0.58, while one of the bioreactors with nitrate feed (bioreactor 12) stagnated after two days of cultivation in batch mode and reached an OD_600_ of 0.23 (**Fig. 3A**). After switching to continuous mode, all three bioreactors with nitrate feed reached similar OD_600_-values of ∼1.2 during the end of Period I, which were 25-30% higher compared to the OD_600_ of the bioreactors with ammonium feed (**Table 1, Fig. 2, Fig. 3A**). The highest observed OD_600_ for the bioreactors with nitrate feed were 1.29 on day 21 during Period IV for bioreactor 10, 1.36 on day 9 during Period II for bioreactor 11, and 1.34 on day 7 during Period I for bioreactor 12 **(Fig. 3, Table 2)**. In comparison, the bioreactors with ammonium feed had the highest average OD_600_-value of 0.90 ± 0.02 on day 4 for Period I (**Fig. 2, Table 1**).

Another noticeable difference was that the OD_600_-values did not remain stable for all bioreactors with nitrate feed during the experiment. Each bioreactor showed fluctuating values at higher OD_600_ before it crashed at different time points during the operating period. For instance, bioreactor 10 underwent a crash in biomass growth on day 8 which continued for the rest of Period II with continuously decreasing OD_600_ (**Fig. 3A**). At this point, the pH stagnated at a pH of 5.8 for bioreactor 10. However, after switching to Period III, bioreactor 10 recovered in OD_600_ back to a value of 1.30 and reached a pH of 5.0 on day 17 of the operating period. During Period IV the OD_600_ was stable for bioreactor 10 (**Fig. 3A**). On day 11 of the operating period, bioreactor 11 underwent a similar crash in biomass growth but, contrarily, did not recover back to high OD_600_-values until the end of the operating period. The pH of this bioreactor 11 remained stable at pH 5.5 from day 11 of the operating period independent of the following periods (**Fig. 3A**). A similar behavior was observed for bioreactor 12, but the abrupt crash in OD_600_ occurred on day 21 during Period III. Bioreactor 12 also did not recover from the crash until the end of the operating period (**Fig. 3A**). It is noteworthy, that we detected nitrate and nitrite in culture samples of bioreactors with nitrate feed undergoing a crash, while neither nitrate nor nitrite were detectable in actively growing or recovering bioreactors with nitrate feed. This indicates a high uptake rate for nitrate by the microbes from the feed medium, and an immediate conversion of the nitrate to ammonium *via* nitrite as an intermediate.

The acetate production rates of all bioreactors with nitrate feed somewhat followed the OD_600_ profile, and reached the highest values that we observed in all our experiments with a maximum value of 129 mmol-C L^−1^ d^−1^ for bioreactor 10 during Period I (**Fig. 3B**). The acetate production rate considerably decreased at the time point of the OD_600_ crashes for the three bioreactors with nitrate feed (**Fig. 3B**). Ethanol production rates were negligible during batch mode for all bioreactors with nitrate feed and slowly increased with decreasing pH during the different periods, after switching to continuous mode, and also considerably dropped after the OD_600_ crashes (**Fig. 3B**). We observed high ethanol production rates in each of the three bioreactors with nitrate feed before the respective OD_600_ crashes with values of 62.0 mmol-C L^−1^ d^−1^ for bioreactor 10, 29.9 mmol-C L^−1^ d^−1^ for bioreactor 11, and 30.6 mmol-C L^−1^ d^−1^ for bioreactor 12 (**Table 2, Fig. 3**). While for bioreactors 11 and 12 the ethanol production rates did not recover after the crashes, for bioreactor 10 the ethanol production rate increased with increasing OD_600_ after the crash, and reached its maximum during Period IV, which is the highest value for the entire study (**Fig. 3B, Table 2**). This value is ∼2.5-fold higher compared to the highest ethanol production rate observed for *C. ljungdahlii* growing with ammonium (**Fig. 2, Table 2**). Acetate production rates, on the other hand, remained low after the recovery of bioreactor 10, which led to the highest measured ethanol/acetate ratio of ∼3.8 with CO_2_ and H_2_ in this study. To our knowledge it is also the highest ethanol/acetate ratio for published studies with acetogens and CO_2_ and H_2_, because Mock et al. (2019) achieved a ratio of ∼1:1. We found that all three bioreactors with nitrate feed behaved differently and underwent stochastic crashes in the OD_600_ at different time points that were most likely connected to a simultaneous accumulation of nitrite. The recovery of one bioreactor from this crash influenced the ethanol and acetate production rates.

Finally, to further test the impact of pH changes on the bioreactors with nitrate feed, we manually adjusted the pH by feeding acid to all bioreactors with nitrate feed to decrease and then keep the pH at pH 4.5 on day 24 of the operating period (**Fig. 3**, indicated with a star symbol). This intervention immediately resulted in a second crash of the OD_600_ and production rates for bioreactor 10, while bioreactors 11 and 12 already were at low OD_600_-values with low acetate and ethanol production rates at that time point.

## Discussion

### Our MBS resulted in reproducible gas-fermentation experiments with *C. ljungdahlii*

The MBS was successfully tested to cultivate *C. ljungdahlii* with CO_2_ and H_2_ in triplicates under various pH conditions during three experiments for four conditions (total of 12 bioreactors). The highly comparable growth behavior of the three replicate bioreactors under batch and continuous conditions for three conditions (*i.e*., Na-acetate, NaCl, and ammonium conditions) confirm a high stability of our MBS (**Fig. S4A, S5A, and S6A)**. We did observe minor differences in the ethanol and acetate production rates between replicates, which were connected to the same medium feed bottle under continuous conditions (**Fig. S4B, S5B, and S6B)**. These differences in single replicates may lead to different production rates, even in controlled bioreactors, and may result from slightly varying gassing or medium feed rates, variations in the pH control, or small but varying diffusion of oxygen into individual bioreactors. This finding clearly indicates the need for replicates during strain characterization and pre-selection in lab-scale bioreactor experiments before scaling up to larger fermentations. With our MBS, we can combine experiments at steady-state conditions for replicates, which safes time in generating statistically relevant data sets. Our future work to further optimize the MBS will target the additional integration of analytic equipment to calculate gas consumption and carbon uptake rates. We had sampled the inlet and outlet gases during all experiments, but our current setup was not sufficient to obtain reliable results. Additional equipment, such as mass-flow controllers, will fill this gap and further increase the data quality during future experiments.

### Acetate feed does not improve ethanol production in *C. ljungdahlii*, but salt stress does at low pH

For our second experiment, we addressed the question whether external acetate feed would increase the ethanol production from CO_2_ and H_2_ for *C. ljungdahlii*, which had been demonstrated for *C. autoethanogenum* (Mock et al., 2015). Both strains are closely related. In contrast to the relatively high ethanol production rates for *C. autoethanogenum*, we observed a negative impact of the acetate feed on growth and production rates of acetate and ethanol (**Results S2, Fig. S4**). The most important differences for our experiment were that for the previous study by Mock et al. (2015): **(1)** a sterile filtration system were used, which allowed the retention of 90% of all microbial cells in the bioreactor; **(2)** a 2-fold higher dilution rate for the medium was applied, which resulted in a 2.3-fold higher feeding rate of acetate; and **(3)** a ∼12-fold higher gas feeding rate of 350 mL min^−1^ was applied. These differences resulted in a 18-fold higher maximum biomass concentration (based on cellular dry weight) for the previous study compared to our experiment (during Period II). Regardless, we had anticipated a higher ethanol production rate for *C. ljungdahlii* with external acetate feed. Therefore, further experiments are required why this did not occur.

For the control, we had supplemented the standard PETC medium with 100 mM NaCl to give a similar ion strength compared to Na-acetate. This resulted in increased ethanol production rates and ethanol/acetate ratios of 0.8 and 1.2 at a lower pH level during Periods III and IV, respectively (**Results S2, Table S2, Fig. S5 and S7**). This is considerably higher than the ethanol/acetate ratios of ∼0.1-0.4 for all other periods in this experiment. While it is likely that the increased salt concentration in the feed medium resulted in a cellular stress response, which has been demonstrated before (Philips et al., 2017), the exact nature of this response in our experiment remains elusive and requires further experimentation. Others have found increased ethanol production rates from syngas with *C. ljungdahlii* in response to cellular stress, which was induced by a low pH in combination with sulfur limitation (Martin et al., 2016), or oxygen exposure (Whitham et al., 2015). The feeding of NaCl at low pH might have interacting effects on ethanol production from CO_2_ and H_2_.

### Feeding nitrate as sole N-source leads to enhanced cell growth even at low pH

For our third experiment, we tested the impact of nitrate as sole N-source on the growth and production rates of acetate and ethanol under pH-controlled conditions. It was recently demonstrated that *C. ljungdahlii* can use nitrate simultaneously for the generation of ammonium (assimilatory nitrate reduction) (Nagarajan et al., 2013), and as an alternative electron acceptor (dissimilatory nitrate reduction) (Emerson et al., 2019). This resulted in enhanced cell growth with sugars or CO_2_ and H_2_ (Emerson et al., 2019). From these findings and our own preliminary batch experiments (**Fig. S8**), we also expected enhanced cell growth in our bioreactor experiment. Our data confirm that the use of nitrate as sole N-source is enhancing CO_2_ and H_2_-dependent growth of *C. ljungdahlii* by up to 50% (based on OD_600_) in continuous mode **(Fig. 3, Table 2)**. Emerson et al. (2019) observed 42% increased growth rates for bottles experiments with CO_2_ and H_2_, while the pH increased from 6.0 to 8.0. We observed a similar increase in the pH-value and a 200% increased final OD_600_ in our preliminary bottle experiments (**Fig. S8**). Our bioreactors with nitrate feed had high OD_600_-values at low pH values, whereas all our other bioreactors showed a correlation between low pH and low OD_600_. We had not anticipated this observation, since acetate production is becoming thermodynamically limited at lower pH (Richter et al., 2016). Consequently, less acetate is produced from acetyl-CoA and, in turn, less ATP is available for the Wood-Ljungdahl pathway (Schuchmann and Müller, 2014). One possible explanation for this observation is that the depleting pool of ATP at low pH is refilled with ATP generated through the reduction of nitrate and concomitant redirection of reducing equivalents. This ATP can then be used for biomass formation.

Our data show that the highest OD_600_ in our bioreactors with nitrate feed remained between an OD_600_ of 1.29 and an OD_600_ of 1.36 during different periods (**Table 2**). This indicates that ATP was not the limiting factor for growth for the bioreactors with nitrate feed and nitrate reduction was sufficient to regenerate redox cofactors. Ethanol formation was not observed in our bottle experiments and in the experiments by Emerson et al. (2019). This led to the hypothesis by the authors that *C. ljungdahlii* predominantly shifts electrons into nitrate reduction rather than towards ethanol formation. The generated ammonium is responsible for the increasing pH-value. In contrast, we demonstrated for all bioreactors with nitrate feed that ethanol production is still possible under pH-controlled conditions (**Fig. 3**). We believe that ethanol formation was absent in our bottle experiments due to the increasing pH-value from ammonia. This is proof that observations with bottles should be followed up with pH-controlled bioreactors.

### Nitrite accumulation indicated a metabolic crash of *C. ljungdahlii* followed by a high ethanol production selectivity after recovering

All three bioreactors with nitrate feed showed different performance behavior during the continuous mode. All these bioreactors underwent a crash at different time points during the cultivation. These crashes were stochastic, because we had already observed the reproducible nature of our MBS. For each of the three bioreactors, we measured an accumulation of nitrite and nitrate at the time point when the crash occurred and afterwards. Before the crashes, we were not able to detect nitrate in any sample. Therefore, we assume that the applied nitrate feed rate of 0.11 mmol h^−1^ (18.7 mM x 0.10 mL min^−1^) was lower than the metabolic uptake rate for nitrate of *C. ljungdahlii*. Our results indicate that an accumulation of nitrite and nitrate is harmful to the microbes and leads to an abrupt halt of the metabolism for yet unknown reasons. A complete physiological characterization of the nitrate metabolism of *C. ljungdahlii*, or any other acetogen, is still missing in literature. Emerson et al. (2019) described that once the applied nitrate was depleted, the culture halted acetate production and crashed (as measured by the OD_600_). The authors explained the crash through an abrupt end of the ATP supply, which is critical to maintain high cell densities for *C. autoethanogenum* (Valgepea et al., 2017). However, the bottle cultures of Emerson et al. (2019) did not crash completely. The OD_600_ decreased by 50% but recovered after a short lag phase, indicating that the remaining CO_2_ and H_2_ was further consumed. An accumulation of nitrite was neither observed during the crash in these experiments nor in our own preliminary bottle experiments (**Fig. S8**). One explanation might be that the metabolic crash was triggered by an insufficient regeneration of NADH. *C. ljungdahlii* possesses two putative hydroxylamine reductases (CLJU_c22260, CLJU_c07730), which could catalyze the reduction of nitrite to ammonium with electrons from NADH (Köpke et al., 2010; Nagarajan et al., 2013). Since we observed simultaneous nitrite and nitrate accumulation in crashing cultures, a metabolic bottleneck at this catalytic step is possible. Another explanation might be that nitrite and/or nitrate inhibit one or several enzymes in *C. ljungdahlii*. Then, as soon as some nitrite and/or nitrate accumulated and inhibited the metabolism, a feedback loop was triggered that quickly led to a complete crash of the metabolism.

For recovering the culture, we assume that the inhibiting compounds have to be washed out of the system to a certain critical threshold. In addition, some removal of the inhibiting compounds due to the recovering activity of the culture would also contribute. For bioreactor 10, we observed a constant decrease of the OD_600_ and acetate and ethanol production rates after the crash in Period I. However, on day 13-14 the decrease started to reach a valley, which indicates that the microbial growth was able to catch up with the dilution of our continuous process. For this bioreactor, the pH in Period II was still high enough to support sufficient growth, and after ∼2 HRT periods the growth rate of the microbes exceeded the dilution rate again, which is indicated by an increasing OD_600_ in the bioreactor (**Fig. 3A**). The recovery of this bioreactor 10 in growth as well as in acetate and ethanol production rates indicates that: (**1)** the nitrate reduction pathway is not *per se* inhibited at low pH; and (**2)** the reduction of nitrate and the production of ethanol is possible simultaneously, and that most likely the low pH triggers a thermodynamic shift towards ethanol production (Richter et al., 2016). However, it remains elusive why the ethanol/acetate ratio in bioreactor 10 reached a nearly ten-fold higher value after recovering from the crash in the presence of nitrate compared to the bioreactors with ammonium feed **(Fig. 3B, Table 2)**.

In contrast, the crash occurred for bioreactor 11 in Period II. While the OD_600_ immediately decreased after the crash, the acetate and ethanol production rates remained somewhat constant until the switch to Period III. However, this bioreactor never recovered from the crash in terms of OD_600_. We believe that the lower pH levels during Period III for bioreactor 11 prevented the growth recovery, which for bioreactor 10 took place at the higher pH level of Period II. We had found reduced growth conditions for all other bioreactors at the lower pH levels (**Fig. 3, Fig. S4**). The same findings hold true for bioreactor 12 for which the growth crash occurred even later. What could be the reason for the stochastic crashes? Valgepea et al. (2017) discussed occurring “crash and recover cycles” during syngas fermentation with *C. autoethanogenum*. They hypothesized that the Wood-Ljungdahl pathway becomes the limiting factor during a period of ample supply of acetyl-CoA at higher biomass and acetate concentration. This can result in an insufficient supply of reducing equivalents due to a loss of H_2_ uptake when the Wood-Ljungdahl pathway cannot keep up anymore. Consequently, the cells are not able to deliver the ATP demand, resulting in a crash. The cells recovered, however, once the extracellular acetate concentration went below a certain threshold, but crashed again after exceeding the threshold. Unfortunately, these threshold acetate concentrations were not given. We observed higher acetate production rates for bioreactor 10 and 11 before the crash compared to those of the bioreactors with ammonium feed (**Fig. 2B, Fig. 3B**). Bioreactor 12 did not reach a similarly high acetate concentration, but the crash occurred at the beginning of Period IV at the lower pH of 4.5. Intrinsically, the extracellular undissociated acetic acid concentration would be higher as a key to trigger the crash event. However, more work is necessary to ascertain the mechanisms of the stochastic crashes.

Nitrate reduction offers a great potential to further optimize gas fermentation of *C. ljungdahlii*. Because ATP limitation is one of the highest burdens to overcome for acetogens (Schuchmann and Müller, 2014; Molitor et al., 2017), the surplus of ATP derived from nitrate reduction could be used to extent the product portfolio towards energy-intense products (Emerson et al., 2019). However, our work clearly demonstrates that nitrate metabolism of *C. ljungdahlii* needs further investigation on both a physiological and a bioprocessing level. The stochastic metabolic crashes demonstrate the importance of replicated bioreactor experiments in the field of acetogen research.

## Acknowledgments

This work was funded through the Alexander von Humboldt Foundation in the framework of the Alexander von Humboldt Professorship, which was awarded to LTA. We are also thankful for additional funding to LTA and BM from the Deutsche Forschungsgemeinschaft (DFG, German Research Foundation) under Germany’s Excellence Strategy – EXC 2124 – 390838134. Finally, we acknowledge support by the DFG and Open Access Publishing Fund of University of Tübingen.

## Author Contributions

CK and LTA designed the MBS. LTA, CK, and BM planned the experiments. CK and NK built, maintained, and sampled the bioreactors. LTA and BM supervised the project. CK analyzed the raw data and drafted the manuscript. All authors edited the manuscript and approved the final version.

## Contribution to the Field Statement

Microbial gas-fermentation with CO_2_ offers great potential to contribute to a climate-friendly and economically feasible production of bio-based chemicals. Various laboratories work on microbial strain design to expand this production platform, while others focus on process engineering in single optimized bioreactors. Consequently, a gap exists between both fields and strain design is often not studied beyond the serum bottle experiment level. We present an open-source multiple-bioreactor system (MBS) for replicable gas-fermentation experiments at a small lab-scale, which fills this gap and provides a transition between strain design and bioprocessing. We show the functionality of our MBS by investigating different physiological changes for the CO_2_-utilizing bacterium *Clostridium ljungdahlii*. Our findings contribute to a better understanding of the microbe in a pH-controlled environment, which is required for bioreactors in biotechnological applications. This will help to look beyond the serum bottle experiment level and enables pre-selection of microbes for further scaling experiments. Furthermore, using our MBS will save time in generating statistically relevant data sets, which is especially relevant for academia.

