## Supplemental Information 1 for "An open-source multiple-bioreactor system for replicable gas-fermentation experiments: Nitrate feed results in stochastic inhibition events, but improves ethanol production of *Clostridium ljungdahlii* with CO_2_ and H_2_"

### *Supplementary Material*

**Includes:**

Results S1 to S2

Figure S1 to S7

Tables S1 to S2

Supplementary PDF-file for the MBS frame

#### Results S1. Concept and assembly of the MBS

All required materials for the MBS were purchased from different manufacturers in Germany or France. We provide a list with all information for each unit of the MBS including manufacturer's names, required amounts, and expected costs (**Table S1**). The basis for our MBS concept was a commercially available 1-L double-walled glass bottle with GLS 80 neck as the bioreactor vessel. The neck of the bottle was additionally flattened by a glass blower to increase the surface contact area between glass and lid. Furthermore, the custom-made lid was sealed with an additional O-ring. We designed a custom-made lid **(Figure S1)**, because it had to offer as many ports as possible to attach pH-regulation and all lines required for a continuous gas and medium feed-in and -out, and we could not find commercially available options that fulfilled our requirements. In addition, the lid had to be autoclavable and gas tight. Our customized lid is made of PTFE and provides vertically attached connection ports for five GL14 fittings and one GL25 fitting. Before attaching the lid, vacuum grease was applied to the glass thread of the bioreactor. One water bath thermostat maintained the temperature of all bioreactors. The equal distribution of the water through black rubber tubing was achieved by installing a starfish manifold. The water outlets of the individual bioreactors were merged into a single line before entering the thermostat again. Stirring bars were used for continuous and unitary agitation of the medium with a multi stirring plate for six bioreactor vessels. Autoclavable pH-electrodes with integrated temperature sensors were installed at the GLS25 port, which was sealed with a PTFE ring that contained a GL25 screw cap. A multi parameter controller, which includes four internal pH/temperature modules, two external pH/temperature modules, and two external relay controllers, was used to track the temperature and to control the pH. All wiring and connections were installed according to the manufacturer's instructions. Twelve peristaltic mini pumps were added to the MBS system and connected to the multi parameter controller to control the pH with base and acid feed for all six bioreactors **(Figure S2)**. A three-part stainless-steel tubing set: a sampling tube, an off-gas tube, and an inlet-gas tube with attached micro sparger was designed, custom-built, and attached to each bioreactor. In addition, the sampling tube was extended by a stainless-steel three-way valve. All GLS14 ports of the bioreactor lid were sealed with 3-part GL14 caps containing PTFE/ETFE replacement inner parts. To maintain continuous feed-in and feed-out conditions, a multichannel pump head equipped with twelve channels was attached to a peristaltic pump. For the mini pumps, a chemical-resistant rubber tubing was used. Furthermore, the multichannel pump head was equipped with 2-stop rubber tubing. All rubber parts were connected *via* Luer-Lock adapters and autoclaved *prior* to use. However, the 2-stop rubber tubing was sterilized with bleach (10 vol-%) and rinsed with sterile water. To keep the feed bottles anaerobic, holes for tubing were drilled through GL45 butyl stoppers. Each feed bottle contained three feed lines, which were connected to the 2-stop rubber tubing of the multichannel pump, one tubing line to add sterile vitamins and reducing agents, and one line (made of rubber tubing and a 10-cm piece of a 1-mL glass pipette). For the gas-out line, rubber tubing coming from the bioreactor went first through a 100-mL serum bottle, which was used as water trap before ending in a fermentation airlock for each bioreactor. The effluent of all bioreactors was collected in a single 10-L bottle.

**Results S2. Growth, acetate, and ethanol production of *C. ljungdahlii* with $CO_2$ and $H_2$ in acetate- or NaCl-supplemented PETC medium using the MBS**

During batch mode, biomass ($OD_{600}$) in all six bioreactors increased to values of 0.59 ± 0.04 (Na-acetate) and 0.55 ± 0.01 (NaCl) (**Figure S4A**). After switching to continuous mode, the $OD_{600}$ increased further to values of 0.80 ± 0.09 (Na-acetate) and 0.76 ± 0.04 (NaCl) on day 6 of Period I (**Figure S4A**). After day 6, the $OD_{600}$ for all bioreactors decreased during Period I and kept decreasing after switching to Period II to values of 0.30 ± 0.08 (Na-acetate) and 0.33 ± 0.06 (NaCl) on day 11 of the experiment (**Figure S4A**). After day 11, the $OD_{600}$ recovered again until the end of Period II, however, for the bioreactors with Na-acetate feed to a lower average value of 0.43 ± 0.08 compared to the bioreactors with NaCl feed with an average value of 0.62 ± 0.04 (**Figure S4A, Table S2**). Noteworthy, the recovery of one of the bioreactors with Na-acetate feed (bioreactor 2) was slower during Period II compared to the other two bioreactors **(Figure S5A)**. However, at the end of this Period, the $OD_{600}$ was similar in all three bioreactors again. During Period III, a decrease of the $OD_{600}$ was observed for the bioreactors with Na-acetate feed to an average value of 0.17 ± 0.04, which continued during Period IV to an almost complete wash-out of cells with an average $OD_{600}$ value of 0.03 ± 0.01 (**Figure S4A, Table S2**). For the bioreactors with NaCl feed we observed a stabilization of the biomass at an average value of 0.55 ± 0.03 during Period III, and a further decrease of the $OD_{600}$ until the end of Period IV with an average $OD_{600}$ of 0.38 ± 0.04 (**Figure S4A**).

During batch mode, the acetate production rates increased with increasing $OD_{600}$, but then considerably decreased after switching to continuous mode (**Figure S4B**). After this decrease, the acetate production rates for all bioreactors increased again to average values of 61.1 ± 0.8 mmol-C $L^{-1}$ $d^{-1}$ (Na-acetate) and 71.8 ± 5.9 mmol-C $L^{-1}$ $d^{-1}$ (NaCl) for Period I (**Figure S4B, Table S2**). For Period II, stable average values of 38.6 ± 1.2 mmol-C $L^{-1}$ $d^{-1}$ (Na-acetate) and 53.1 ± 2.9 mmol-C $L^{-1}$ $d^{-1}$ (NaCl) were observed (**Figure S4B, Table S2**). While for the bioreactors with NaCl feed the average acetate production rates were 40.1 ± 1.7 mmol-C $L^{-1}$ $d^{-1}$ and 27.1 ± 1.4 mmol-C $L^{-1}$ $d^{-1}$ during Periods III and IV, respectively, the average acetate production rates for the bioreactors with Na-acetate feed decreased to 20.3 ± 4.6 mmol-C $L^{-1}$ $d^{-1}$ (Period III), and as low as 2.2 ± 1.5 mmol-C $L^{-1}$ $d^{-1}$ (Period IV). Ethanol production rates were negligible during batch mode, but slowly increased to average values of 6.7 ± 1.0 mmol-C $L^{-1}$ $d^{-1}$ (Na-acetate) and 4.2 ± 0.5 mmol-C $L^{-1}$ $d^{-1}$ (NaCl), and 13.0 ± 0.8 mmol-C $L^{-1}$ $d^{-1}$ (Na-acetate) and 10.6 ± 1.5 mmol-C $L^{-1}$ $d^{-1}$ (NaCl) in Period I and II, respectively (**Figure S4B, Table S2**). While for the bioreactors with Na-acetate feed the ethanol production rates decreased to 4.7 ± 1.2 mmol-C $L^{-1}$ $d^{-1}$ (Period III) and even to no ethanol production (Period IV), the ethanol production rate of the bioreactors with NaCl feed increased further to 32.0 ± 1.0 mmol-C $L^{-1}$ $d^{-1}$ (Period III) and 31.8 ± 2.5 mmol-C $L^{-1}$ $d^{-1}$ (Period IV) (**Figure S4B, Table S2**).

The feeding of NaCl in bioreactors 7/8/9 resulted in an 8-fold higher NaCl concentration compared to standard PETC medium. In these bioreactors with nitrate feed during the Period III and IV, the ethanol production rates exceeded those of the bioreactors with ammonium feed by ~2-fold and ~8-fold, respectively (**Table 1, Figure 2B, Table S2, Figure S3B**). Furthermore, while the ethanol production rates for the bioreactors with nitrate feed were constantly high at ~32 mmol-C $L^{-1}$ $d^{-1}$ during Period III and IV, the $OD_{600}$ was 35% lower during Period IV compared to Period III (**Table S2, Figure S3A**). This resulted in an increased ethanol/acetate ratio (based on production rates) of 0.8 during Period III, and 1.2 during Period IV (**Table S2**), compared to 0.3 and 0.1 for the bioreactors with ammonium feed (**Table 2**).

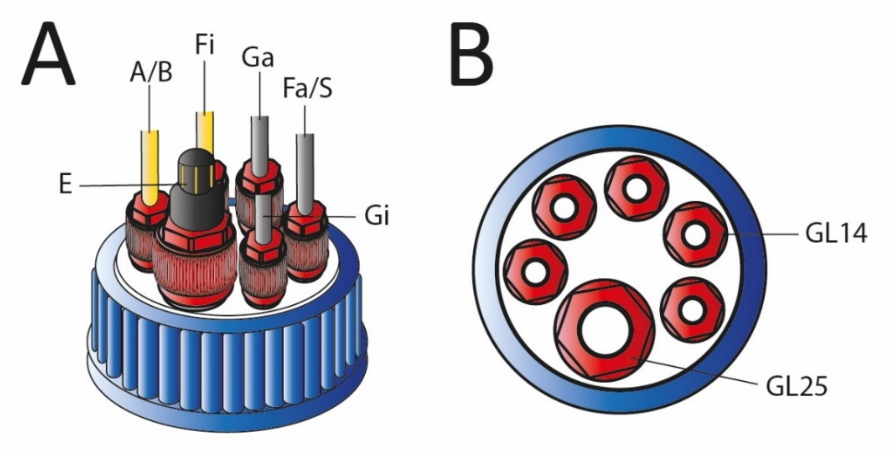


**Figure S1. Side and top view of the customized lid designed for the MBS.** Side view on the lid with connected tubing, ports, and pH-electrode (A). Top view on the lid surface (B). The lid meets all requirements to perform either heterotrophic or autotrophic cultivation experiments with attached pH-control and continuous medium feed-in and -out. Abbreviations: E, pH-electrode; A/B, acid or/and base feed; Fi, medium feed-in line; Ga, gas-out line; Fa/S, medium feed-out line/sampling port *via* three-way valve; Gi, gas inlet, GL14, thread size GL14; GL25, thread size GL25.

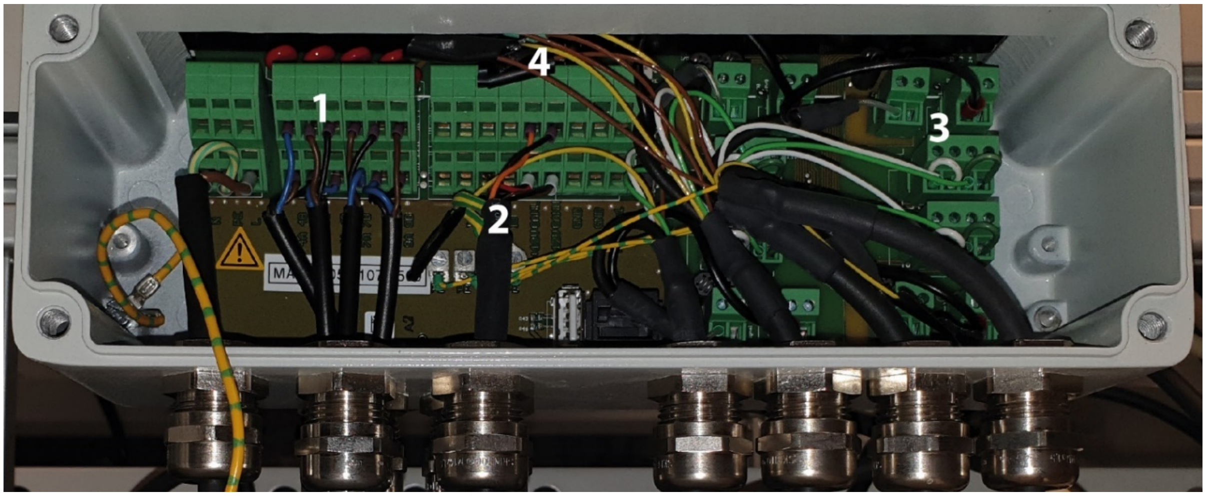


**Figure S2. Wiring of the multi parameter controller KM3000 to operate six pH-electrodes and twelve pumps *via* relay and CAN bus interface.** Four internal relay signals are used to control four pumps directly through 3-wire cables (1). The other eight pumps are controlled as in-line signal *via* CAN bus (2). Four internal interfaces are used to connect four pH/Pt-1000 electrodes (3). It is important to have small bridge connectors for the white and the green wire (not provided with the cable combinations). The other two pH/Pt-1000 electrodes are connected in-line *via* CAN bus. The yellow and brown wire are not required and insulated for safety reasons (4). Unless otherwise stated, all connections were made according to the manufacturer's instructions.

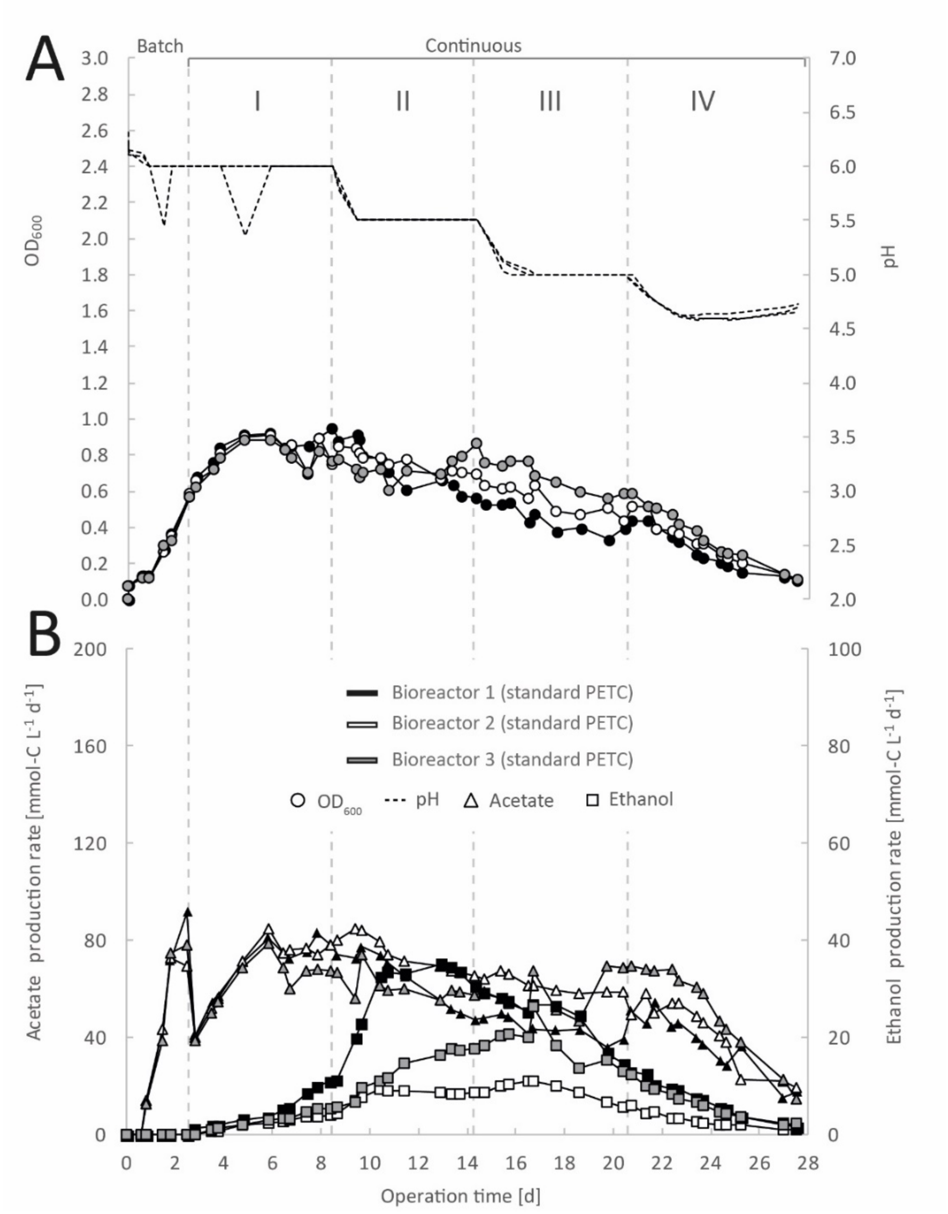


**Figure S3. Single bioreactor data for continuous gas fermentation of *C. ljungdahlii* with $CO_2$ and $H_2$ in standard PETC medium at different periods.** Single values for pH and $OD_{600}$ (A), and for acetate and ethanol production rates in mmol-C $L^{-1}$ $d^{-1}$ (B). The horizontal dotted lines indicate the continuous mode in which medium with different pH was fed to each bioreactor. The pH was not regulated with feeding acid in continuous mode, but by adjusting the feed medium pH to the desired value and by biological acetic acid production. The cultivation volume was initially 500 mL but was tracked daily during the experiment ranging from 500-650 mL. The gas feed rate was 30 mL $min^{-1}$. The medium feed was 0.10 mL $min^{-1}$. The bioreactors were operated at 37°C and 300 rpm for 27.5 days. Period: I, pH=6.0; II, pH=5.5, III, pH=5.0; and IV, pH=4.5.

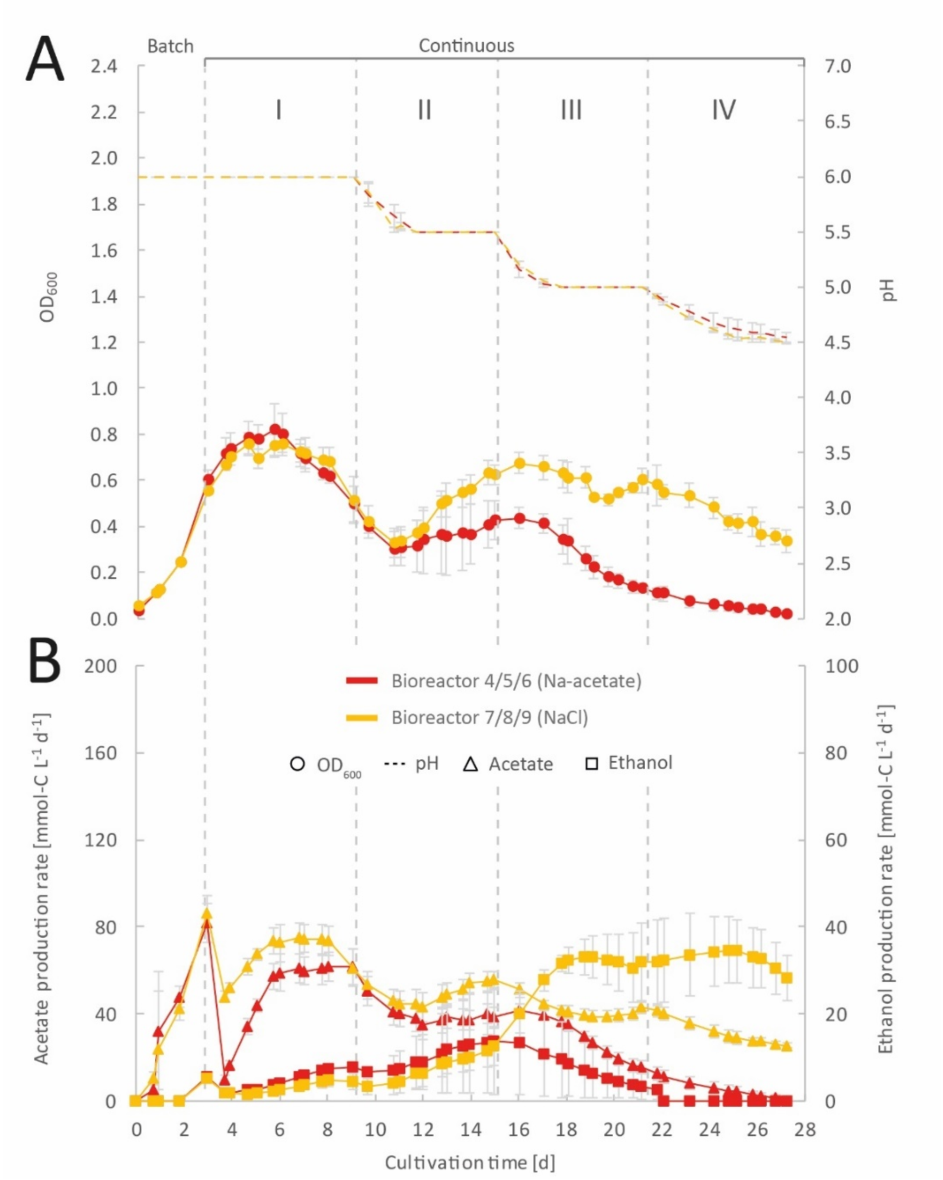


**Figure S4. Impact of feeding acetate to continuous gas fermentation of *C. ljungdahlii* with $CO_2$ and $H_2$ at different periods.** Mean values of triplicates with standard deviation (n=3) for pH and $OD_{600}$ (A), and for acetate and ethanol production rates in mmol-C $L^{-1}$ $d^{-1}$ (B). The bioreactors with Na-acetate feed were fed with medium that contained 100 mM Na-acetate (red). The bioreactors with NaCl feed were fed with medium that contained 100 mM NaCl (yellow). The horizontal dotted lines indicate the continuous mode in which medium with different pH was fed to each bioreactor. The pH was not regulated with feeding acid in continuous mode, but by adjusting the feed medium pH to the desired value and by biological acetic acid production. The cultivation volume was initially 500 mL but was on average 600 mL during continuous mode. The gas feed rate was 30 mL $min^{-1}$. The medium feed rate was 0.10 mL $min^{-1}$. The fed medium was adjusted to the desired pH before autoclaving. The bioreactors were operated at 37°C and 300 rpm for 27.2 d. Period: I, pH=6.0; II, pH=5.5, III, pH=5.0; and IV, pH=4.5.

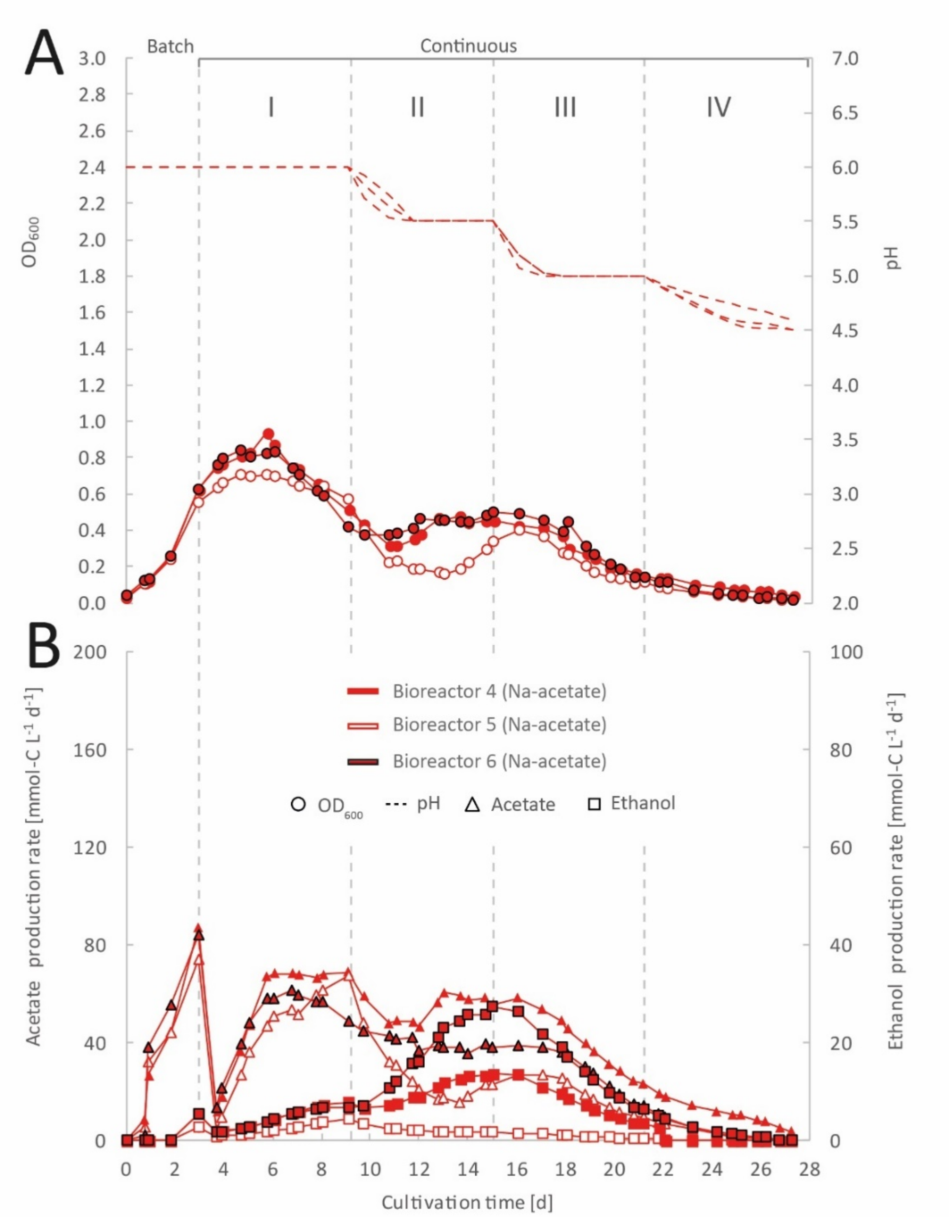


**Figure S5. Single bioreactor data describing the impact of feeding acetate to continuous gas fermentation of *C. ljungdahlii* with $CO_2$ and $H_2$ at different periods.** Single values for pH and $OD_{600}$ (A), and for acetate and ethanol production rates in mmol-C $L^{-1}$ $d^{-1}$ (B). The bioreactors with Na-acetate feed were fed with medium that contained 100 mM Na-acetate. The horizontal dotted lines indicate the continuous mode in which medium with different pH was fed to each bioreactor. The pH was not regulated with feeding acid in continuous mode, but by adjusting the feed medium pH to the desired value and by biological acetic acid production. The cultivation volume was initially 500 mL but was on average 600 mL during continuous mode. The gas feed rate was 30 mL $min^{-1}$. The medium feed was 0.10 mL $min^{-1}$. The fed medium was adjusted to the desired pH before autoclaving. The bioreactors were operated at 37°C and 300 rpm for 27.2 d. Period: I, pH=6.0; II, pH=5.5, III, pH=5.0; and IV, pH=4.5.

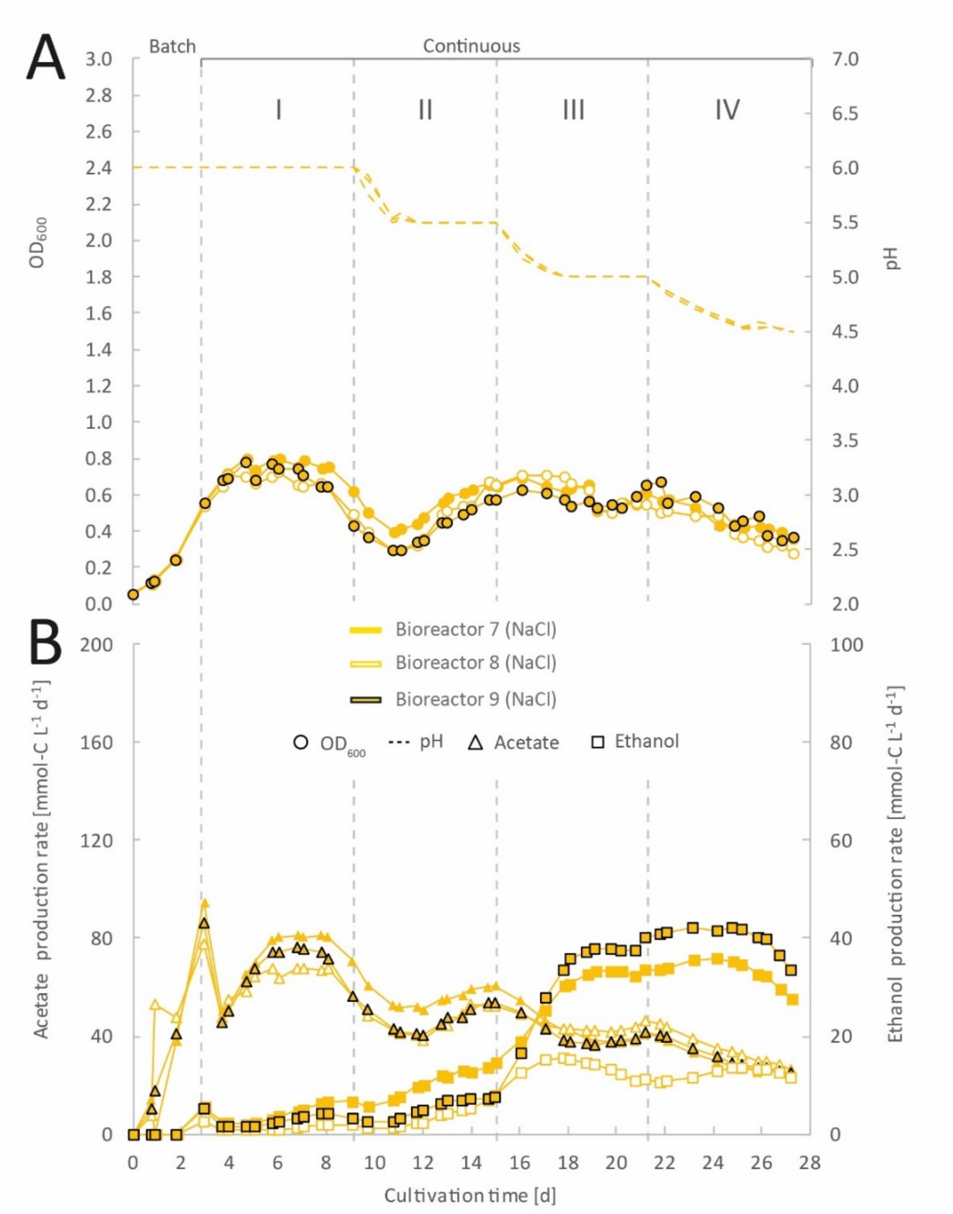


**Figure S6. Single bioreactor data describing the impact of feeding additional NaCl to continuous gas fermentation of *C. ljungdahlii* with $CO_2$ and $H_2$ at different periods.** Single values for pH and $OD_{600}$ (A), and for acetate and ethanol production rates in mmol-C $L^{-1}$ $d^{-1}$ (B). The bioreactors with Na-acetate feed were fed with medium that contained 100 mM NaCl. The horizontal dotted lines indicate the continuous mode in which medium with different pH was fed to each bioreactor. The pH was not regulated with feeding acid in continuous mode, but by adjusting the feed medium pH to the desired value and by biological acetic acid production. The cultivation volume was initially 500 mL but was on average 600 mL during continuous mode. The gas feed rate was 30 mL $min^{-1}$. The medium feed rate was 0.10 mL $min^{-1}$. The fed medium was adjusted to the desired pH before autoclaving. The bioreactors were operated at 37°C and 300 rpm for 27.2 d. Period: I, pH=6.0; II, pH=5.5, III, pH=5.0; and IV, pH=4.5.

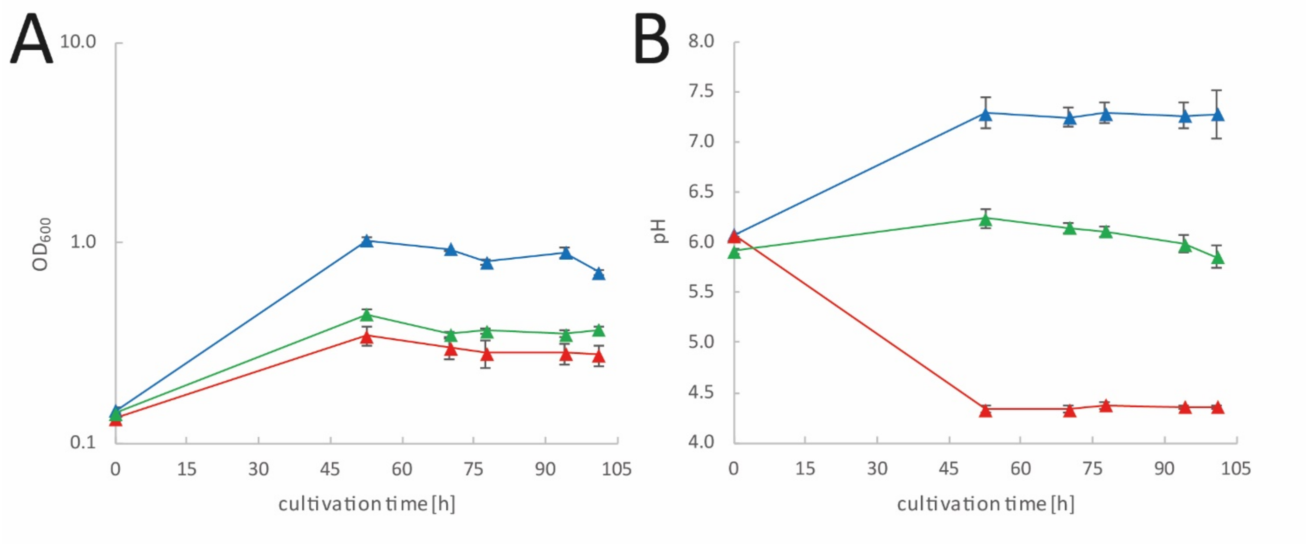


**Figure S7. Bottle cultivation of *C. ljungdahlii* with $CO_2$ and $H_2$ using ammonium, nitrate, or a mixture of both as N-source.** Growth as $OD_{600}$ in triplicate (n=3) under each tested condition (A). Changes in pH (B). 50 mL PETC medium supplemented with 0.5 g/L yeast extract but without fructose and ammonium chloride was provided in 240-mL serum bottles. Na-nitrate, ammonium chloride, or a mixture of both N-sources were added as sterile and anaerobic solution. The bottles were filled with sterile $CO_2$ and $H_2$ gas (20/80 vol-%) to 1 bar overpressure before inoculation. Pre-cultures were grown heterotrophically in PETC with fructose for 48 h at 37°C in a stand incubator, washed once with sterile PBS, concentrated, and directly used for inoculation. Cultivation of main cultures were carried out at 150 rpm in a shaking incubator (Lab companion, ISS-7100R, Jeio Tech, Korea) at 37°C for 101 hours. *C. ljungdahlii* growing with 20 mM Na-nitrate (▲); *C. ljungdahlii* growing with 20 mM ammonium chloride (▲); and *C. ljungdahlii* growing with a mixture of 10 mM Na-nitrate and 10 mM ammonium chloride (▲).

**Table S1. Required materials for the MBS to operate six bioreactors.** The list comprises details about every item and summarizes the total cost of the system (as of Spring, 2019).

| Item | Manufacturer | Item no. | Quantity | Approx. cost in €* |
|---|---|---|---|---|
| 1-L double-walled bottle | Duran/VWR | 215-4157 | 6 | 2000 |
| Customized bioreactor lid | Bohlender/Zinnstag | XZ019-182117 | 6 | 2000 |
| O-ring (70x3 mm, FPM30) | Reif | 4141280 | 6 | 25 |
| Multi stirring plate | 2mag | 2MAG_10020 | 1 | 1250 |
| Set of customized stainless-steel tubing, Ø 6mm, tube (320 mm) with sparger, sample tubing (280 mm), off-gas tubing (100 mm) | bbi-biotech | quote (BZV-2018100900491) | 6 | 2250 |
| Gas manifold for 6 lines (self-built) | Swagelock | - | 1 | 1500 |
| KM3000 Multi-Parameter controller, including 4 internal relays + Can-Bus-Interface (standard version) | Xylem/Si-analytics | 90278011 | 1 | 2050 |
| Internal pH/temp module | Xylem/Si-analytics | 90278011 | 4 | 1500 |
| External pH/temp module for KM3000 | Xylem/Si-analytics | 90278011 | 2 | 850 |
| REL 2000 CAN (external module) 4 relays, CAN-Bus-Interface | Xylem/Si-analytics | 90278017 | 2 | 850 |
| pH-electrode (SL 81-225 pHT VP, pt1000) | Xylem/Si-analytics | 90279050 | 6 | 2600 |
| Cable combination for SL 81-225 pHT VP | Xylem/Si-analytics | 85442000 | 6 | 1100 |
| Set of shrink tubing green/yellow (3 mm) | Conrad | 541743-62 | 1 | 5 |
| Set of wire end ferrule 0.25 mm x 6 mm | Conrad | 739539-62 | 1 | 5 |
| Masterflex C/L Mini pumps (13 to 80 rpm) | ColeParmer | 77122-14 | 12 | 9400 |
| Heating thermostat (CC-104A) | Huber | 461-1056 | 1 | 1600 |
| Octagon-Manifold | Interchim | 343938 | 1 | 440 |
| Customized bioreactor frame | Item24 | | 1 | 1200 |
| Plastic 10mm, ESD, grey (300x600mm) | Item24 | 0.0.614.87 | 2 | 15 |
| Plastic 10mm, ESD, grey (300x200mm) | Item24 | 0.0.614.87 | 1 | 5 |
| GL14 cap, 3-parts, including PTFE/ETFE fittings (6 mm) | Bola | D590-06 | 30 | 550 |
| GL25 cap with PTFE ring | Bola | H984-03 + H975-18 | 6 | 60 |
| Magnetic stirring bar (38 mm) | Carl Roth | A954.1 | 6 | 40 |
| Airlock | Chemglass/VWR | AF-0513 | 6 | 350 |
| Water trap (240-mL serum bottle) | Glasgerätebau Ochs | 102.041 | 10 | 20 |
| Masterflex Multichannel pump head + 12 cartridges | ColeParmer | HV-07519-25/SI-07519-85 | 1 | 2900 |
| Masterflex L/S pump (100 RPM) | ColeParmer | HV-07528-30 | 1 | 1600 |
| 2-Stop pump tubing for multichannel pump | ColeParmer | HV-06431-26 | 1 | 150 |
| Masterflex mini pump tubing | ColeParmer | GZ-95809-30 | 6 | 30 |
| Norprene tubing in size 14 | ColeParmer | GZ-06402-14 | 1 | 75 |
| Norprene tubing in size 16 | ColeParmer | GZ-06402-16 | 1 | 75 |
| Masterflex I/P Norprene Tubing A 60 G | ColeParmer | GZ-06404-73 | 1 | 50 |
| Luer/Lock fittings in different sizes | Carl Roth | CT59.1-64.1 | 100 | 150 |
| 3-way valves | Carl Roth | P340.1 | 6 | 200 |
| 5-L duran bottle | Duran/VWR | 215-0057 | 4 | 360 |
| 10-L duran bottle | Duran/VWR | 215-0058 | 1 | 150 |
| Butyl stoppers for GL45 | Glasgerätebau Ochs | 444704 | 4 | 15 |
| Tube clamps | Carl Roth | YE70-1 | 50 | 100 |
| Zip ties and holder for 6 mm | Hornbach (hardware store) | - | 100 | 30 |
| Screws and nuts | Hornbach (hardware store) | - | 20 | 10 |
| * includes 19% tax | | | Total cost | **37560** |

**Table S2. Average values for $OD_{600}$ and production rates during the continuous fermentation of *C. ljungdahlii* with $CO_2$ and $H_2$ at four different periods.**

| Bioreactor | $OD_{600}$[1] | Acetate production rate [mmol-C $L^{-1}$ $d^{-1}$][1] | Ethanol production rate [mmol-C $L^{-1}$ $d^{-1}$][1] | $Ratio_{Et/Ac}$[2] |
|---|---|---|---|---|
| | | **Period I (pH 6.0)** | | |
| Bioreactor 4/5/6 (Na-acetate) | 0.63 ± 0.09 | 61.1 ± 0.8 | 6.7 ± 1.0 | 0.1 |
| Bioreactor 7/8/9 (NaCl) | 0.66 ± 0.08 | 71.8 ± 5.9 | 4.2 ± 0.5 | 0.1 |
| | | **Period II (pH 5.5)** | | |
| Bioreactor 4/5/6 (Na-acetate) | 0.39 ± 0.03 | 38.6 ± 1.2 | 13.0 ± 0.8 | 0.3 |
| Bioreactor 7/8/9 (NaCl) | 0.58 ± 0.05 | 53.1 ± 2.9 | 10.6 ± 1.5 | 0.2 |
| | | **Period III (pH 5.0)** | | |
| Bioreactor 4/5/6 (Na-acetate) | 0.17 ± 0.04 | 20.3 ± 4.6 | 4.7 ± 1.2 | 0.2 |
| Bioreactor 7/8/9 (NaCl) | 0.55 ± 0.03 | 40.1 ± 1.7 | 32.0 ± 1.0 | 0.8 |
| | | **Period IV (pH 4.5)** | | |
| Bioreactor 4/5/6 (Na-acetate) | 0.03 ± 0.01 | 2.2 ± 1.5 | n.d.[3] | 0.0 |
| Bioreactor 7/8/9 (NaCl) | 0.38 ± 0.04 | 27.1 ± 1.4 | 31.8 ± 2.5 | 1.2 |

[1] Values for the bioreactors with Na-acetate and NaCl feed (n=3) are given as the average (± standard deviation) from three bioreactors of the last 5 data points of every period.

[2] Et, Ethanol; Ac, Acetate; [3] n.d., not detected.
