## Supplemental Information 2 for "An open-source multiple-bioreactor system for replicable gas-fermentation experiments: Nitrate feed results in stochastic inhibition events, but improves ethanol production of *Clostridium ljungdahlii* with CO_2_ and H_2_"

### Project overview

Project name: MBS frame blueprint

Project number: 40afc9d6b7ecec6553ffad72df53b7871

i24.ws?40afc9d6b7ecec6553ffad72df53b7871

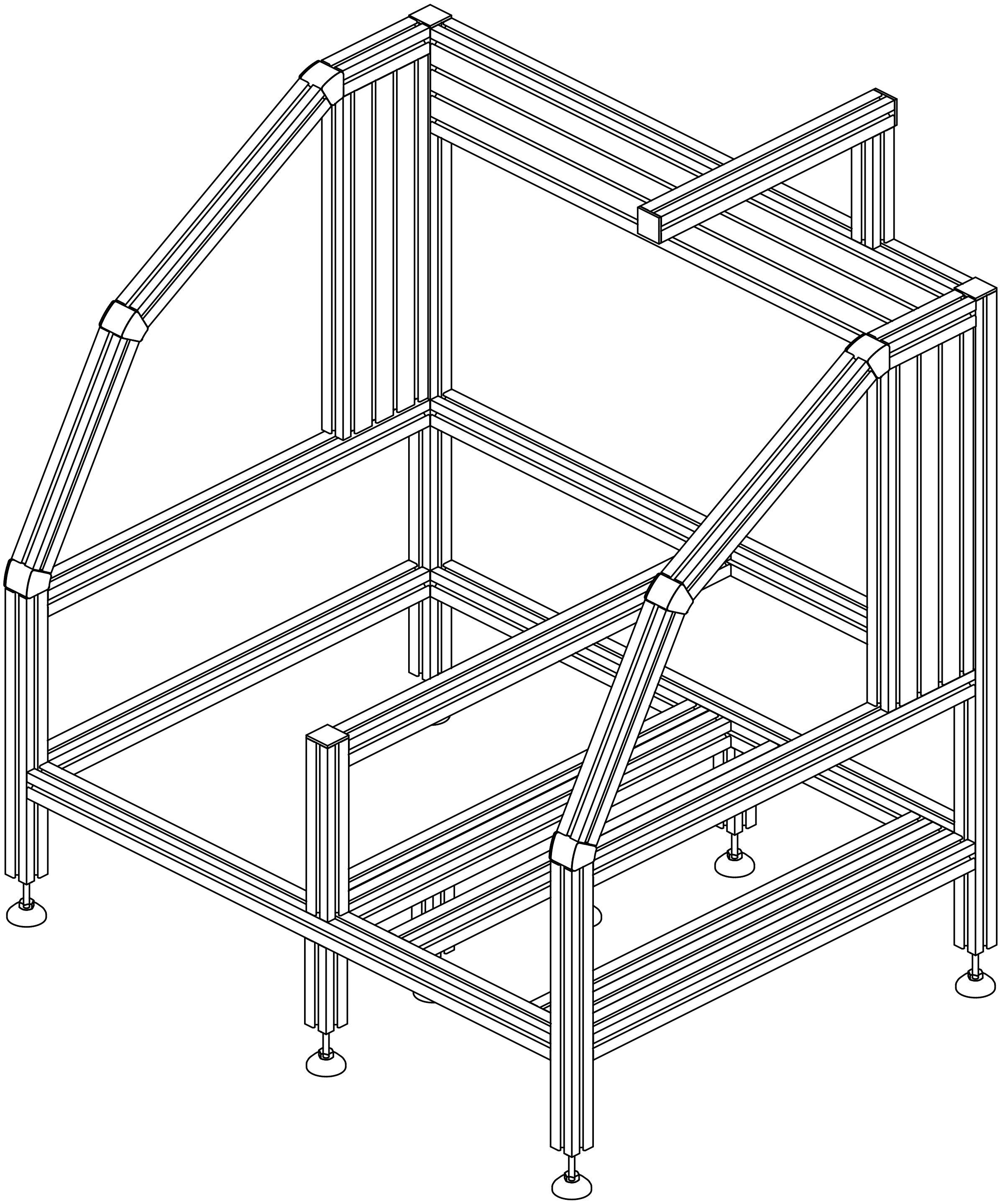

### Assembly overview

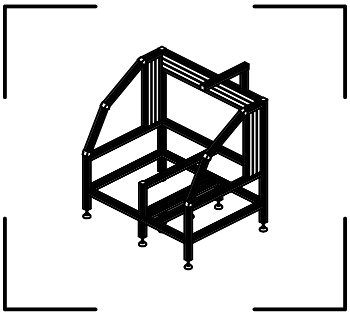

**+2 Separately placed articles**

### Parts list

| Position | Article designation | Article No. | Side | Quantity (All together) | Quantity (Assembly 1) | Quantity (Separately) |
|---|---|---|---|---|---|---|
| 1 | Profile 6 120x30, natural, Length: 396.51mm | 0.0.419.04 | - | 2 | 2 | - |
| 2 | Profile 6 120x30, natural, Length: 740mm | 0.0.419.04 | - | 1 | 1 | - |
| 3v | Profile 6 30x30, natural, Length: 100mm | 0.0.419.01 | 6 | 4 | 4 | - |
| 4 | Profile 6 30x30, natural, Length: 123.7mm | 0.0.419.01 | - | 2 | 2 | - |
| 5 | Profile 6 30x30, natural, Length: 130mm | 0.0.419.01 | - | 1 | 1 | - |
| 6 | Profile 6 30x30, natural, Length: 201mm | 0.0.419.01 | - | 1 | 1 | - |
| 7 | Profile 6 30x30, natural, Length: 250mm | 0.0.419.01 | - | 2 | 2 | - |
| 8 | Profile 6 30x30, natural, Length: 300mm | 0.0.419.01 | - | 2 | 2 | - |
| 9 | Profile 6 30x30, natural, Length: 330mm | 0.0.419.01 | - | 1 | 1 | - |
| 10v | Profile 6 30x30, natural, Length: 330mm | 0.0.419.01 | 8 | 2 | 2 | - |
| 11 | Profile 6 30x30, natural, Length: 540mm | 0.0.419.01 | - | 10 | 8 | 2 |
| 12 | Profile 6 30x30, natural, Length: 740mm | 0.0.419.01 | - | 3 | 3 | - |
| 13v | Profile 6 30x30, natural, Length: 755mm | 0.0.419.01 | 10 | 2 | 2 | - |
| 14 | Profile 6 R30/60-30°, natural, Length: 30mm | 0.0.459.54 | - | 6 | 6 | - |
| 15 | Cap 6 30x30, black | 0.0.419.22 | - | 5 | 5 | - |
| 16 | Cap 6 R30/60-30°, black | 0.0.459.39 | - | 12 | 12 | - |
| 17 | Knuckle Foot D40, M8x80, black | 0.0.265.69 | - | 8 | 8 | - |
| 18 | Automatic-Fastening Set 6, bright zinc-plated | 0.0.419.71 | - | 82 | 82 | - |

#### Fastener Technology (All together)

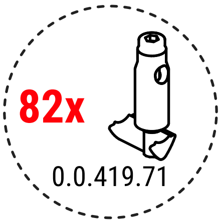

### Unmachined profiles

**2x**

Part 1, Profile 6 120x30, natural
Article No.: 0.0.419.04
Length: 396.51mm, Unmachined

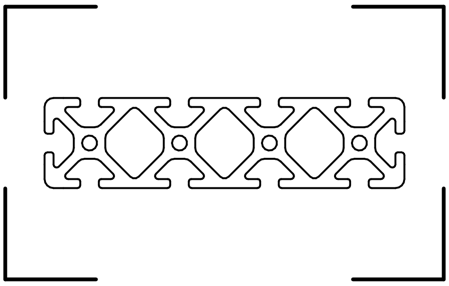

i24.ws?40afc9d6b7ecec6553ffad72df53b7871.1

Part 2, Profile 6 120x30, natural
Article No.: 0.0.419.04
Length: 740mm, Unmachined

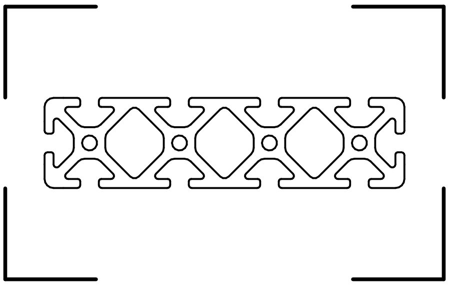

i24.ws?40afc9d6b7ecec6553ffad72df53b7871.2

**2x**

Part 4, Profile 6 30x30, natural
Article No.: 0.0.419.01
Length: 123.696mm, Unmachined

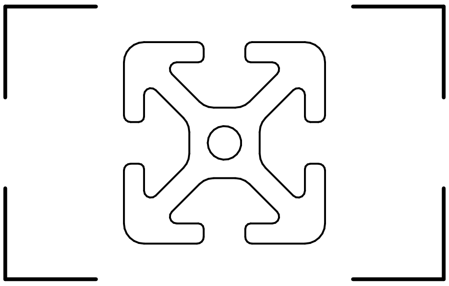

i24.ws?40afc9d6b7ecec6553ffad72df53b7871.4

Part 5, Profile 6 30x30, natural
Article No.: 0.0.419.01
Length: 130mm, Unmachined

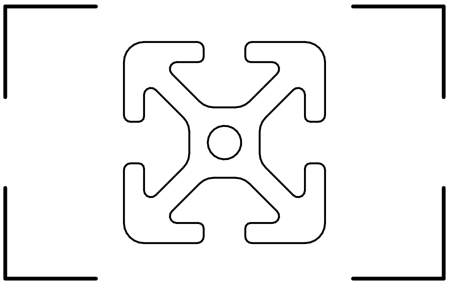

i24.ws?40afc9d6b7ecec6553ffad72df53b7871.5

Part 6, Profile 6 30x30, natural
Article No.: 0.0.419.01
Length: 201mm, Unmachined

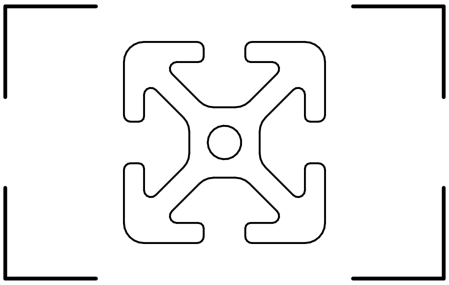

i24.ws?40afc9d6b7ecec6553ffad72df53b7871.6

**2x**

Part 7, Profile 6 30x30, natural
Article No.: 0.0.419.01
Length: 250mm, Unmachined

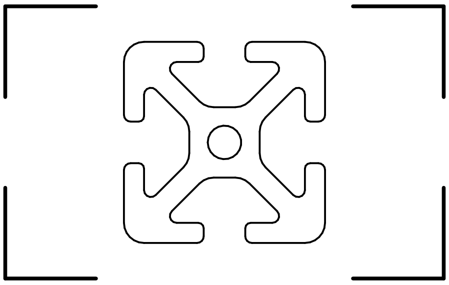

i24.ws?40afc9d6b7ecec6553ffad72df53b7871.7

### Unmachined profiles

**2x**

Part 8, Profile 6 30x30, natural
Article No.: 0.0.419.01
Length: 300mm, Unmachined

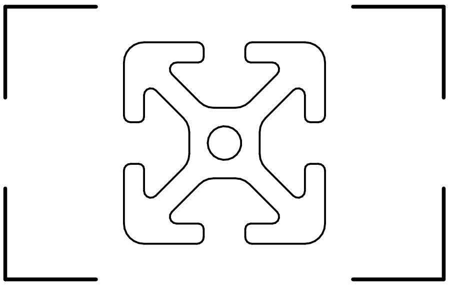

i24.ws?40afc9d6b7ecec6553ffad72df53b7871.8

Part 9, Profile 6 30x30, natural
Article No.: 0.0.419.01
Length: 330mm, Unmachined

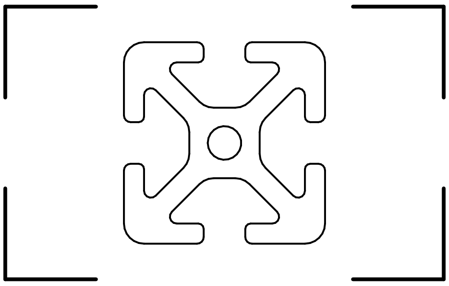

i24.ws?40afc9d6b7ecec6553ffad72df53b7871.9

**10x**

Part 11, Profile 6 30x30, natural
Article No.: 0.0.419.01
Length: 540mm, Unmachined

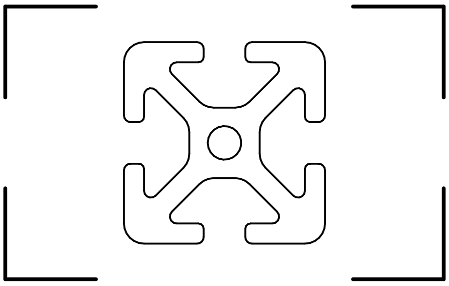

i24.ws?40afc9d6b7ecec6553ffad72df53b7871.11

**3x**

Part 12, Profile 6 30x30, natural
Article No.: 0.0.419.01
Length: 740mm, Unmachined

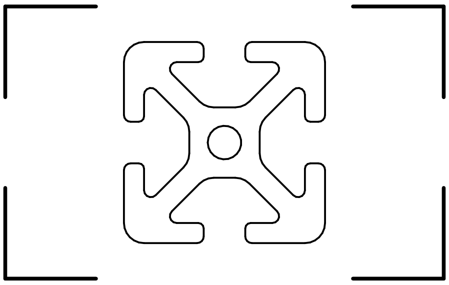

i24.ws?40afc9d6b7ecec6553ffad72df53b7871.12

**6x**

Part 14, Profile 6 R30/60-30°, natural
Article No.: 0.0.459.54
Length: 30mm, Unmachined

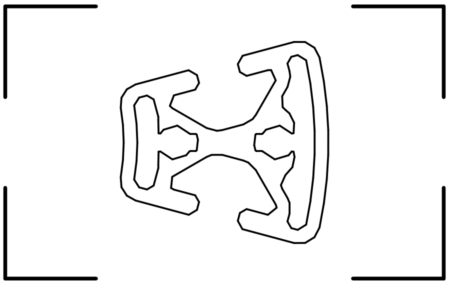

i24.ws?40afc9d6b7ecec6553ffad72df53b7871.14

4x

### Machining processes Part 3v, Profile 6 30x30, natural
#### Article No.: 0.0.419.01, Length: 100mm

View of end face 1

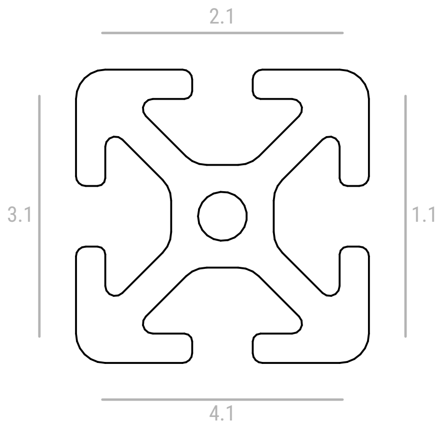

i24.ws?40afc9d6b7ecec6553ffad72df53b7871.3v

| Machining type | Side | Number | Dimension (end face 1) | Machining process designation | Dimension (end face 2) |
|---|---|---|---|---|---|
| D6.8 drilled hole with M8 thread | End face 1 | v1 | - | M8x64 | - |

**End face 1**
Side 2
is at the top

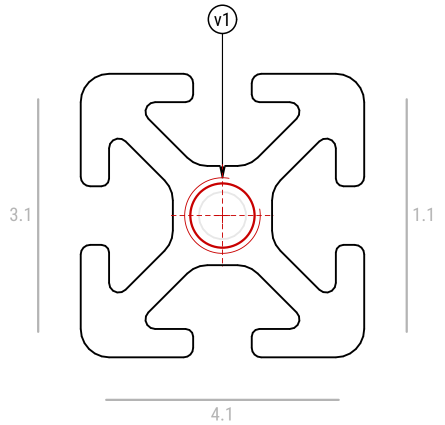

M8x64

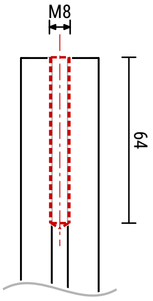

**v1**

### Control view (machining processes), part 3v

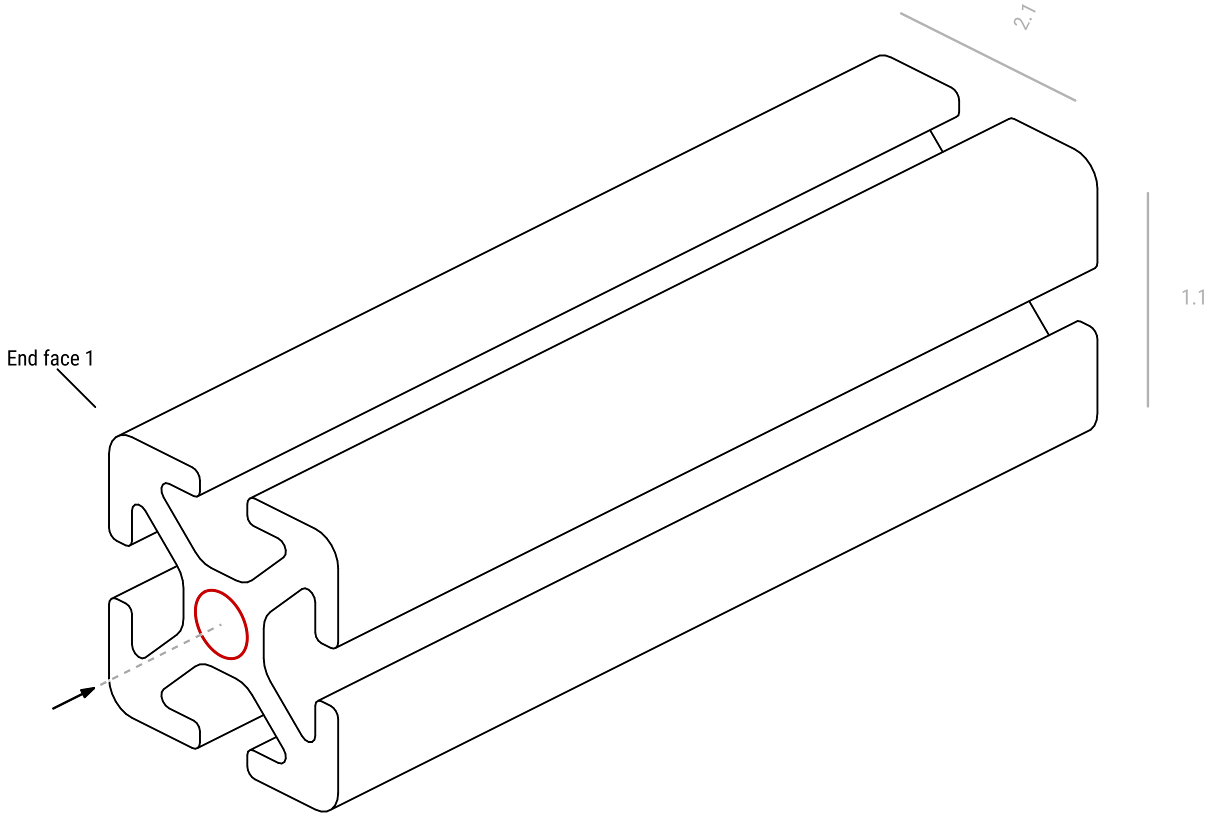

**2x**

### Machining processes Part 10v, Profile 6 30x30, natural
#### Article No.: 0.0.419.01, Length: 330mm

View of end face 1

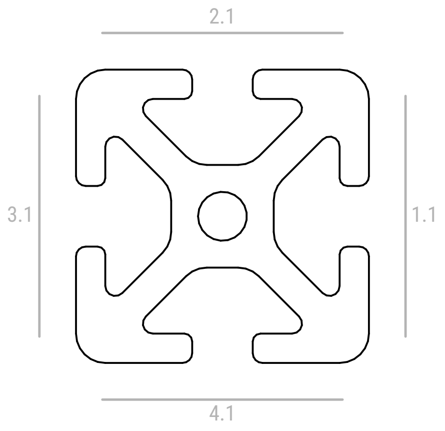

i24.ws?40afc9d6b7ecec6553ffad72df53b7871.10v

| Machining type | Side | Number | Dimension (end face 1) | Machining process designation | Dimension (end face 2) |
|---|---|---|---|---|---|
| D6.8 drilled hole with M8 thread | End face 2 | v1 | - | M8x64 | - |

**End face 2**

Side 2
is at the top

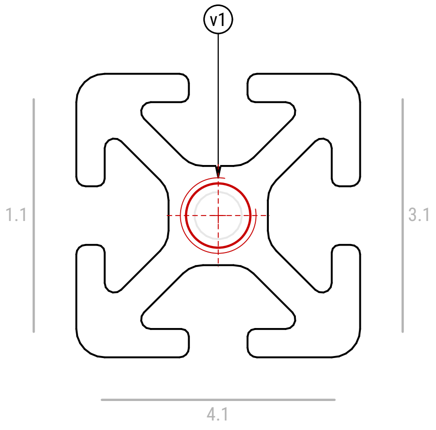

M8x64

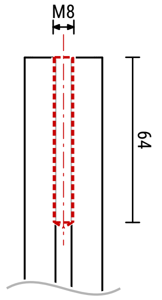

**v1**

### Control view (machining processes), part 10v

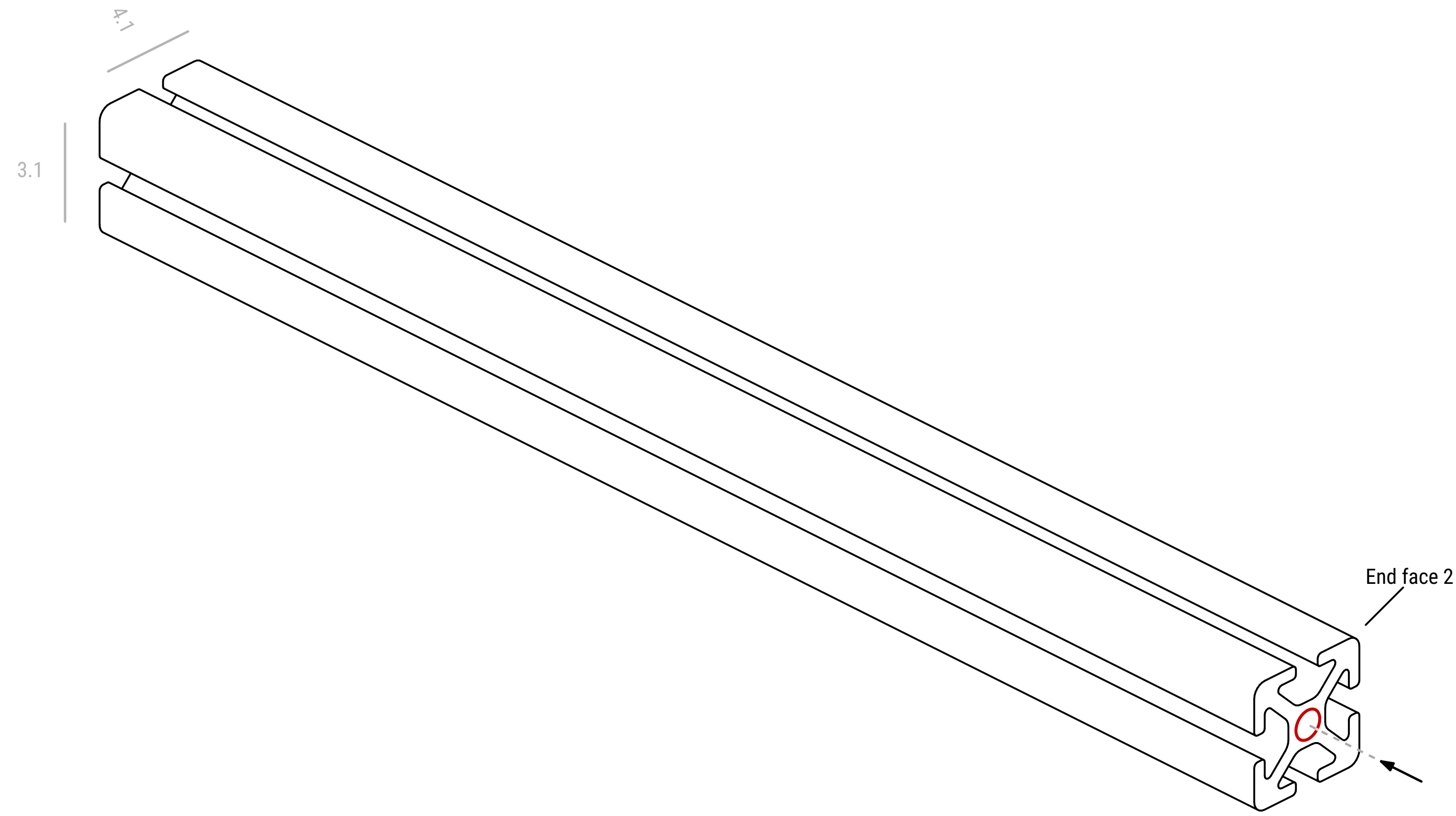

2x

### Machining processes Part 13v, Profile 6 30x30, natural
### Article No.: 0.0.419.01, Length: 755mm

View of end face 1

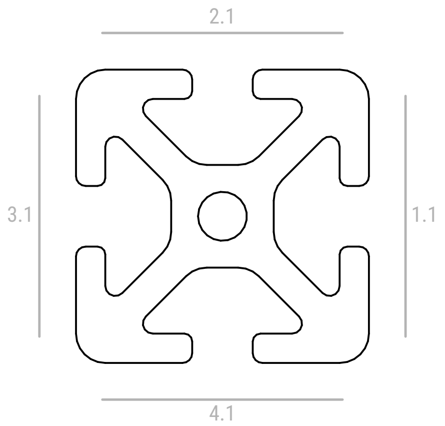

i24.ws?40afc9d6b7ecec6553ffad72df53b7871.13v

| Machining type | Side | Number | Dimension (end face 1) | Machining process designation | Dimension (end face 2) |
|---|---|---|---|---|---|
| D6.8 drilled hole with M8 thread | End face 2 | v1 | - | M8x64 | - |

**End face 2**

Side 2
is at the top

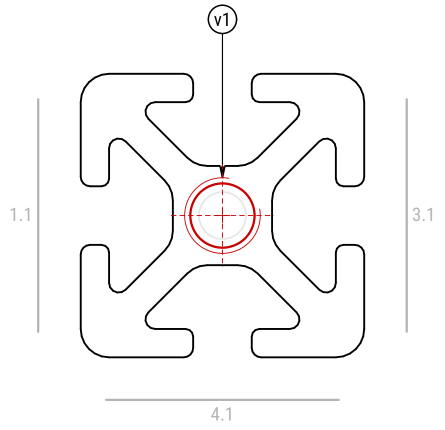

M8x64

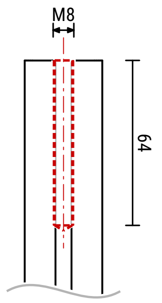

**v1**

### Control view (machining processes), part 13v

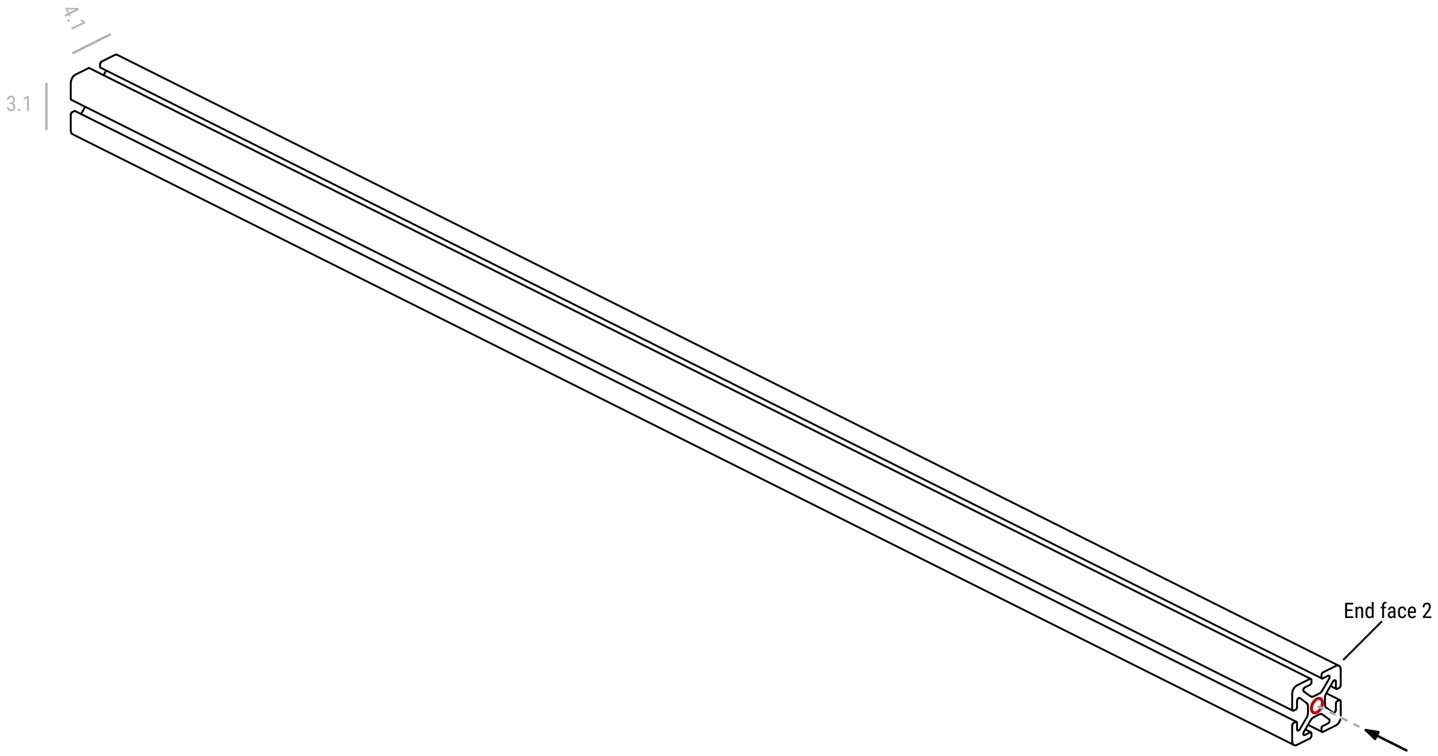

### Isometric view

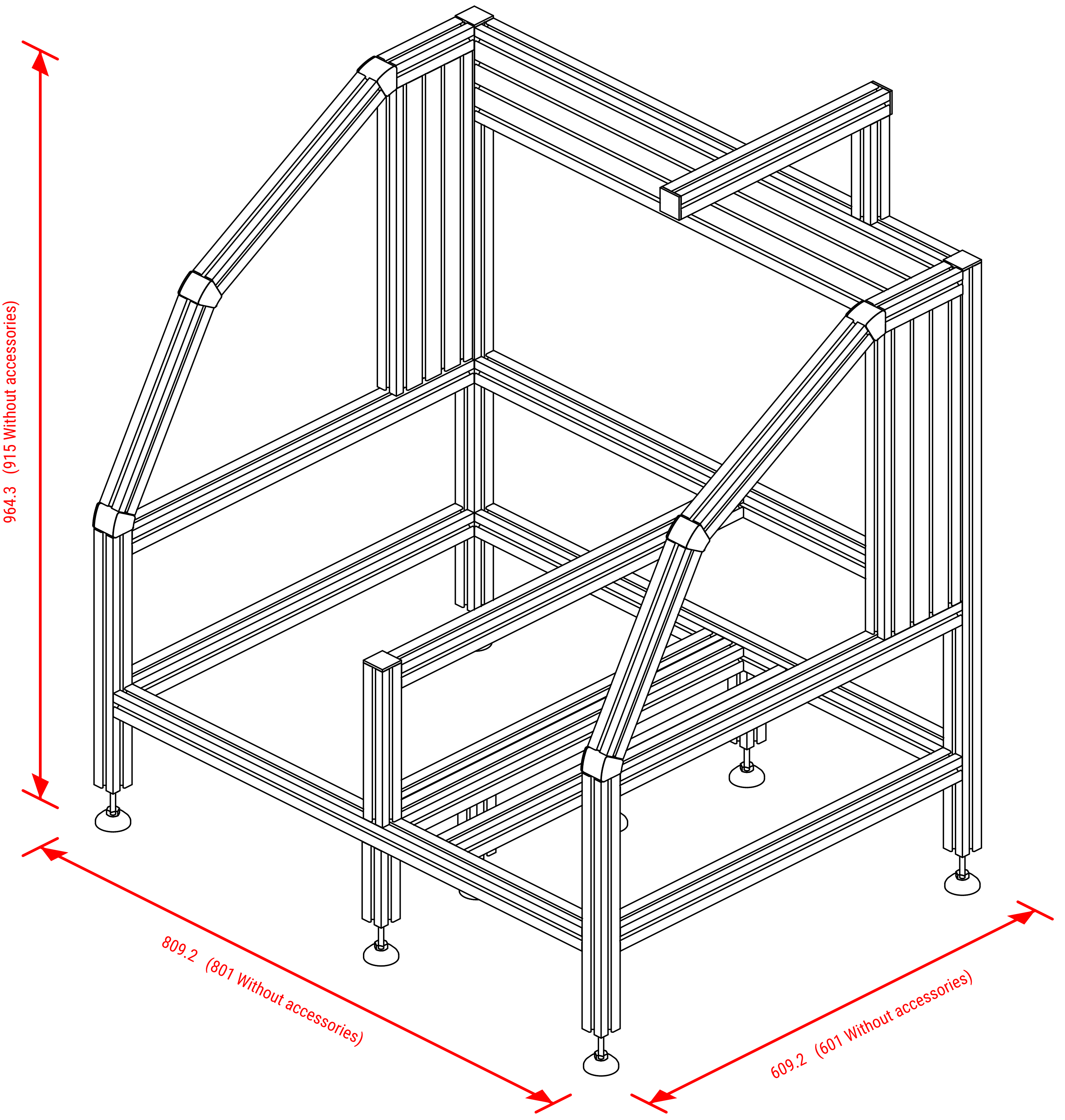

### Multiview projection

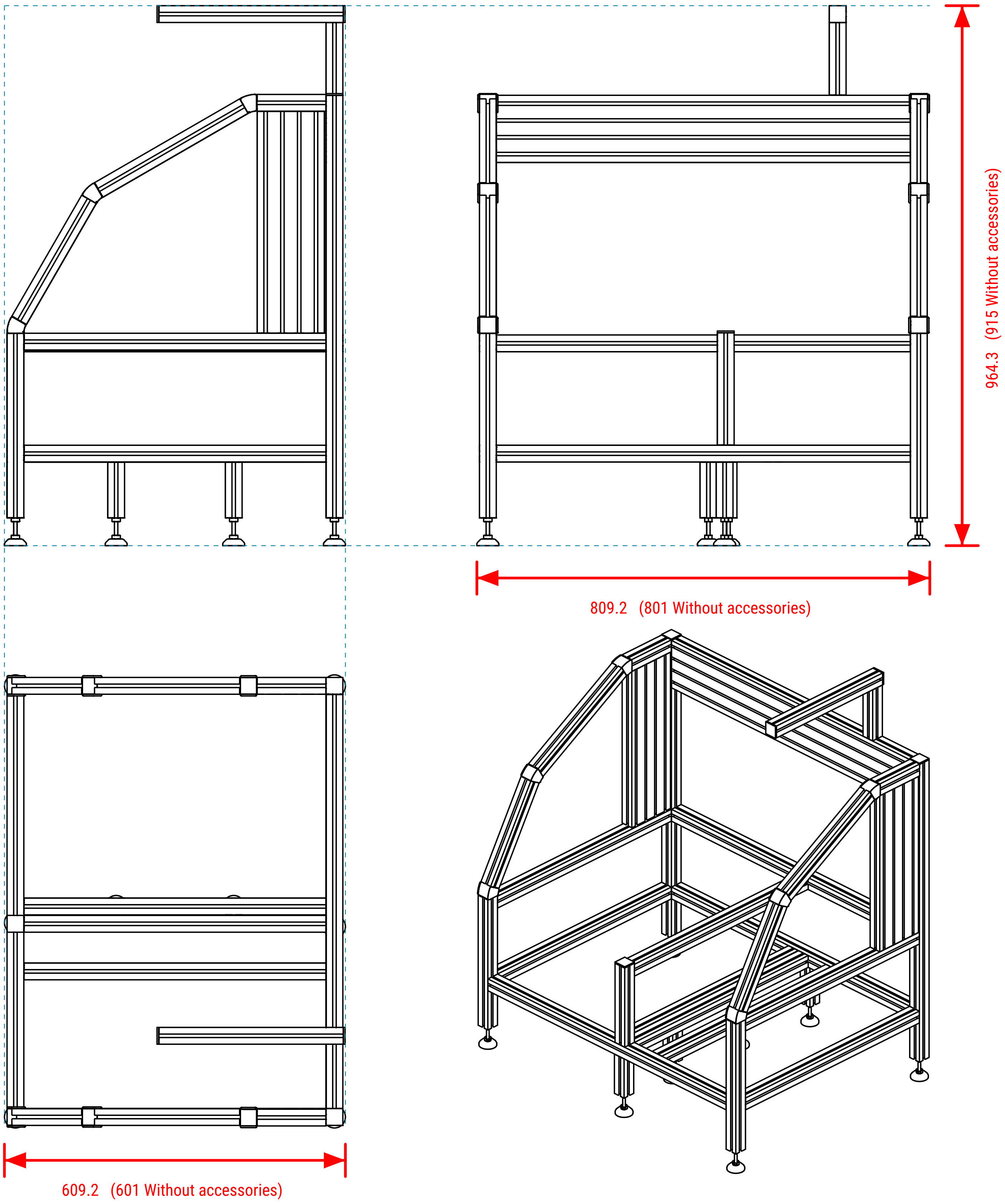

### Exploded view ( Profiles )

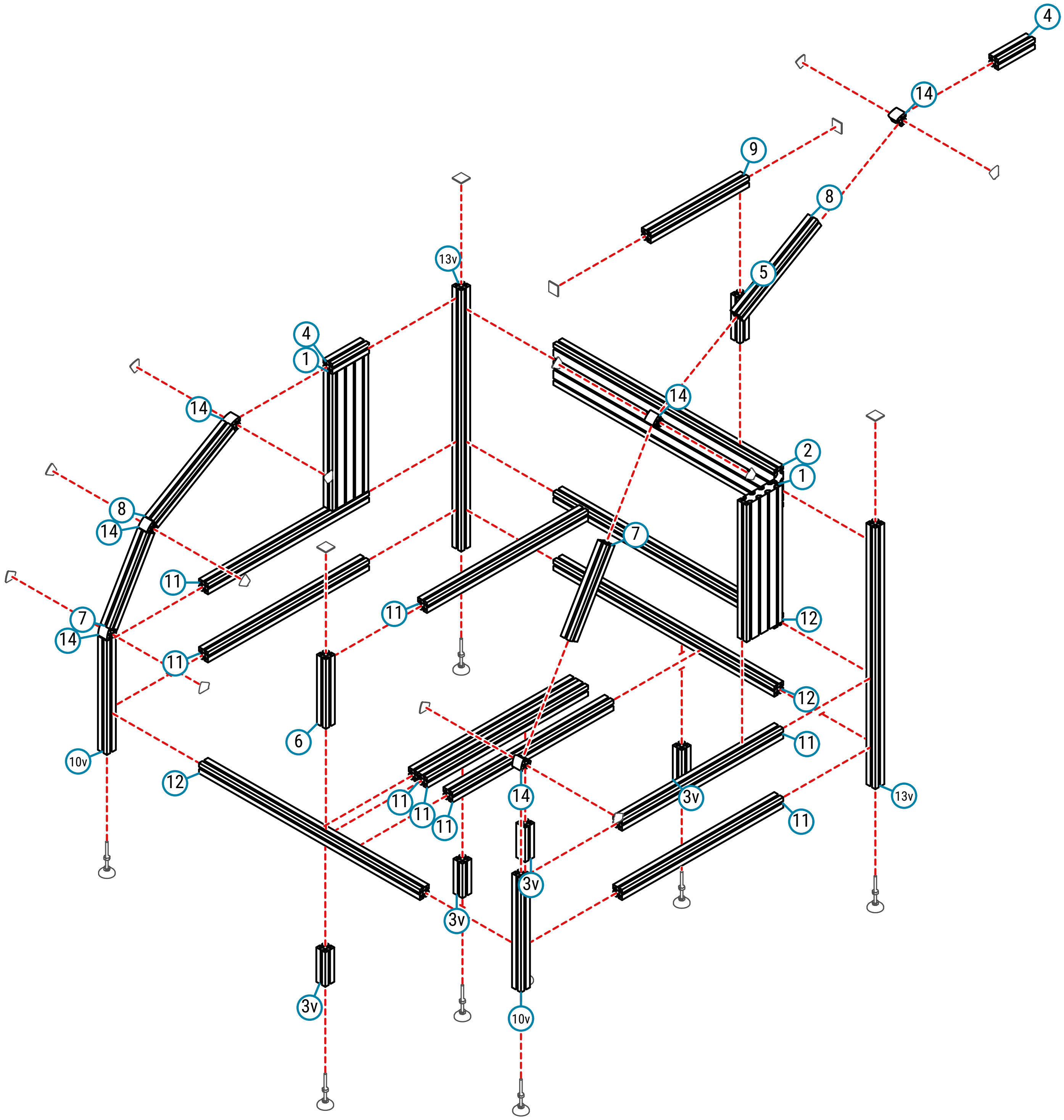

Project : MBS frame blueprint, 40afc9d6b7ecec6553ffad72df53b7871

### Exploded view ( accessories and Fastener Technology )

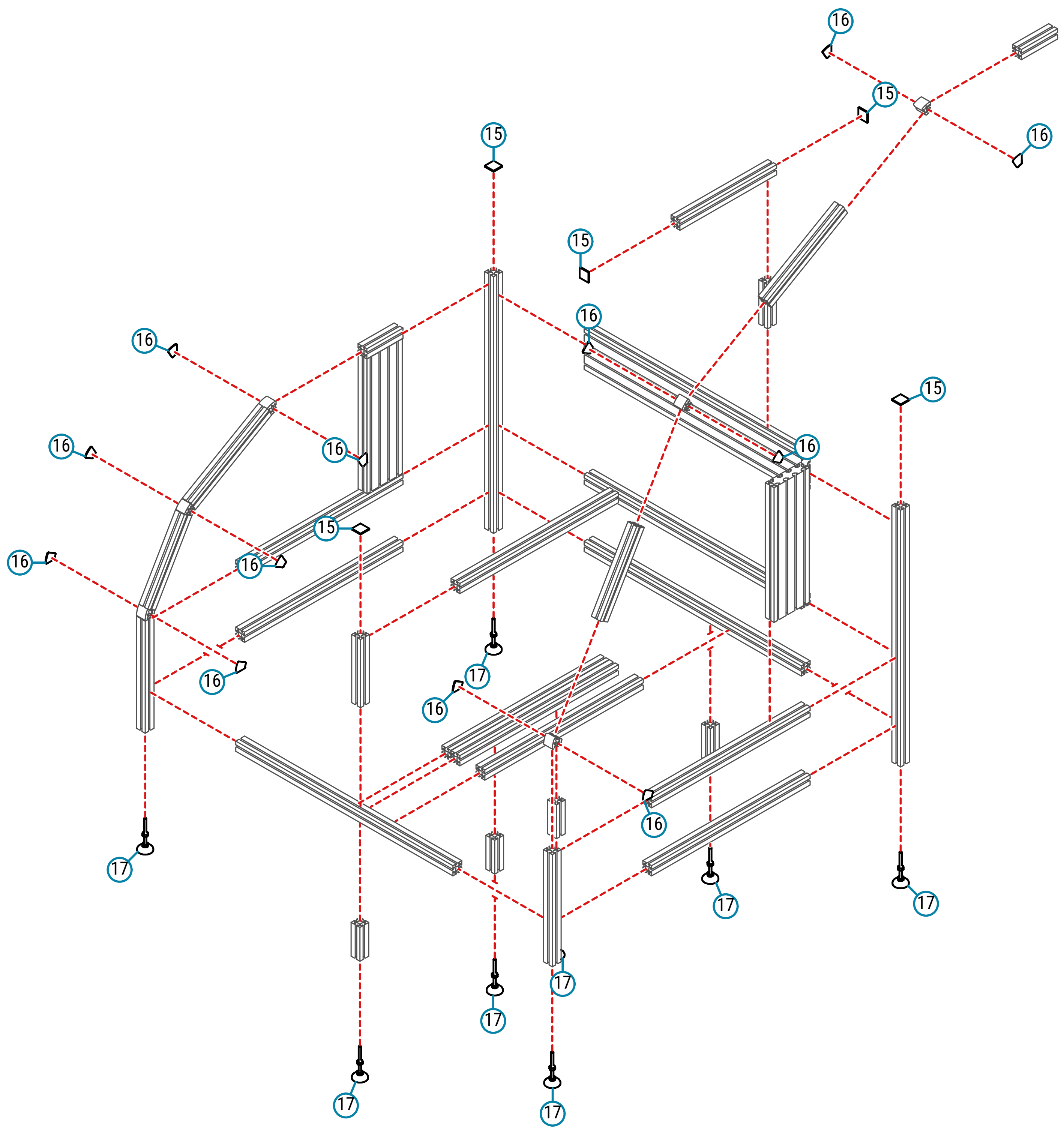

### Installation guide

#### Step 1/ 17

Start with part 16 and 13v

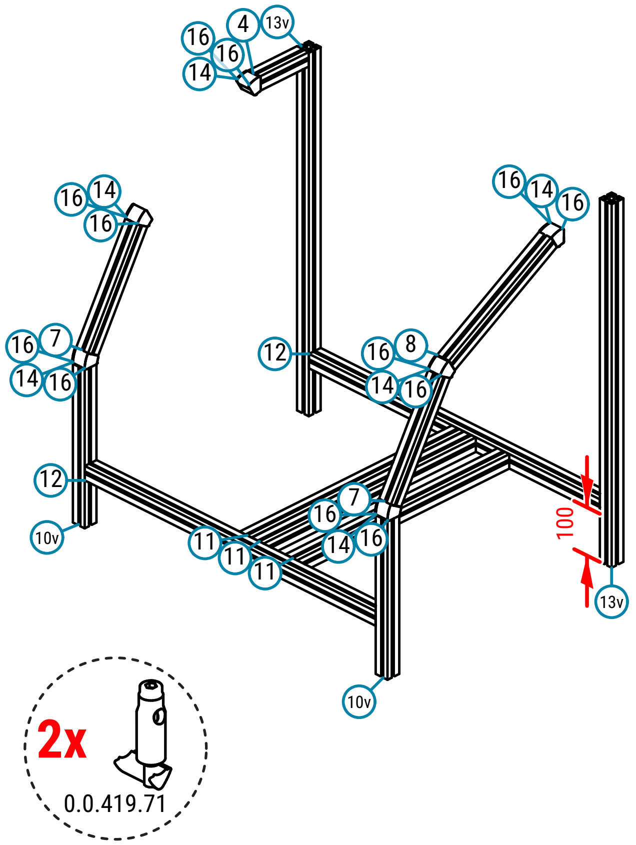

#### Step 2/ 17

Install 1 x part 11

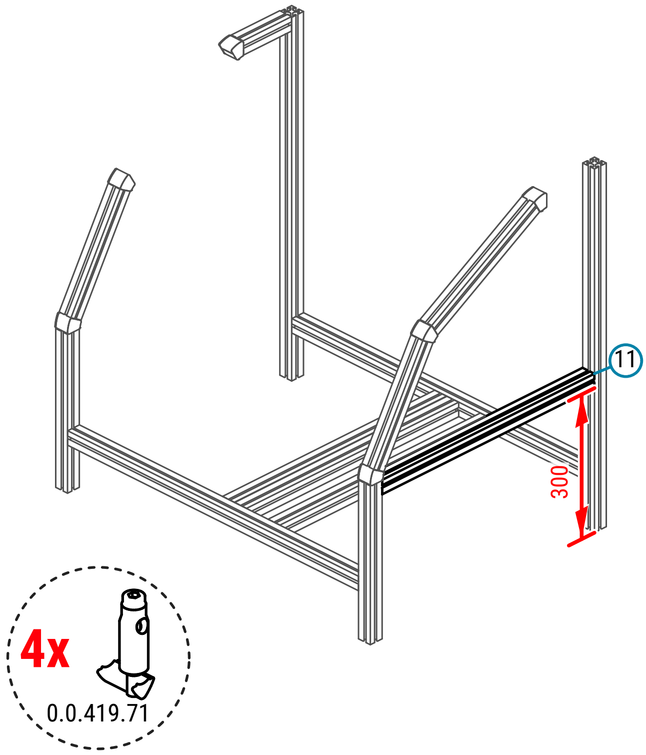

#### Step 3/ 17

Install 1 x part 8

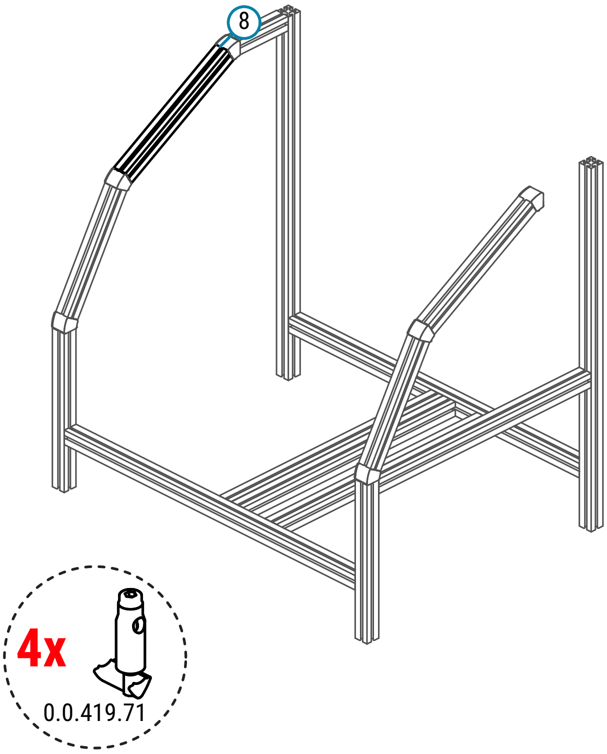

#### Step 4/ 17

Install 1 x part 12

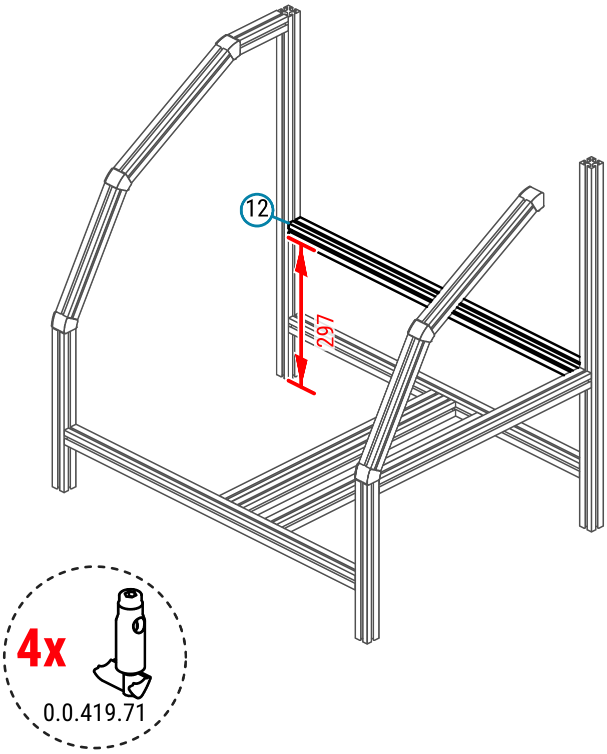

#### Step 5/ 17

Install 1 x part 2

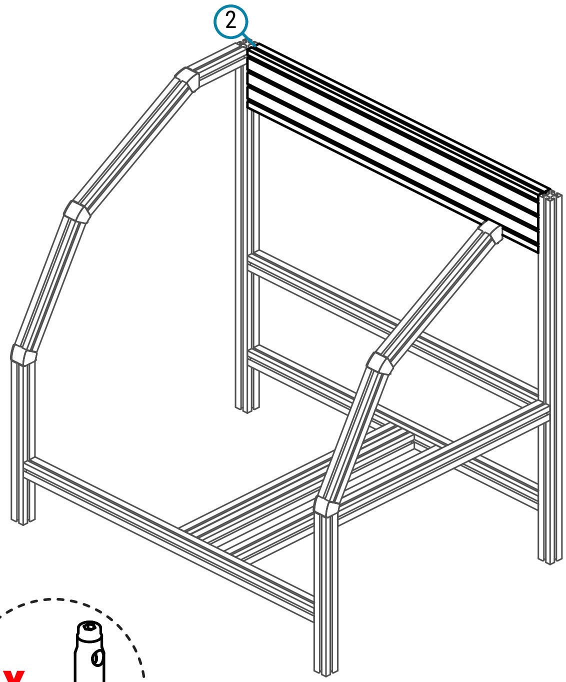

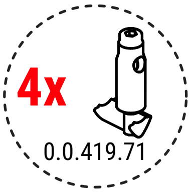

#### Step 6/ 17

Install 1 x part 1

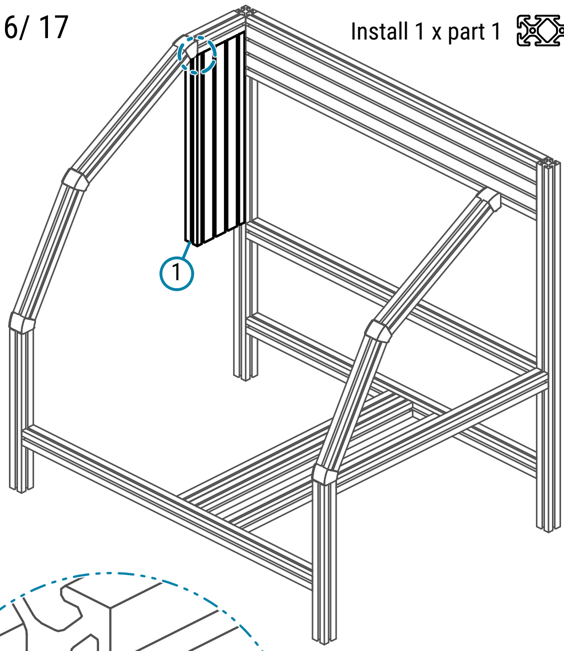

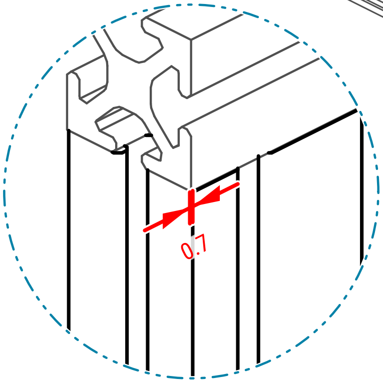

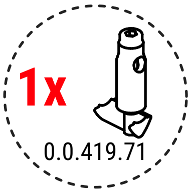

#### Step 7/ 17

Install 2 x part 11

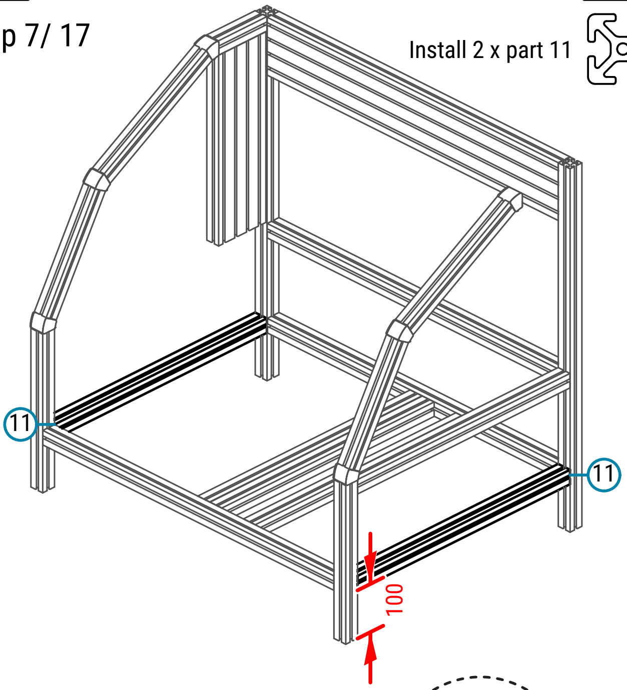

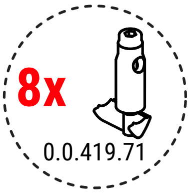

All installed parts have exactly the same dimension

#### Step 8/ 17

Install 1 x part 11

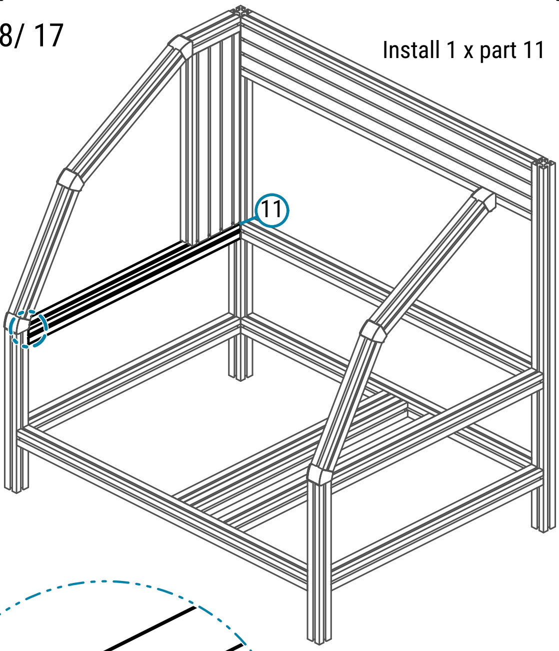

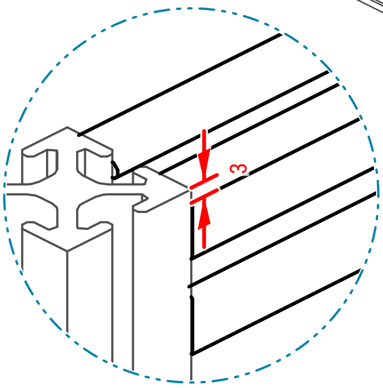

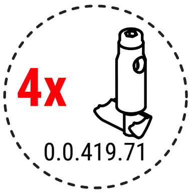

#### Step 9/ 17

Install 1 x part 1

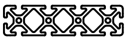

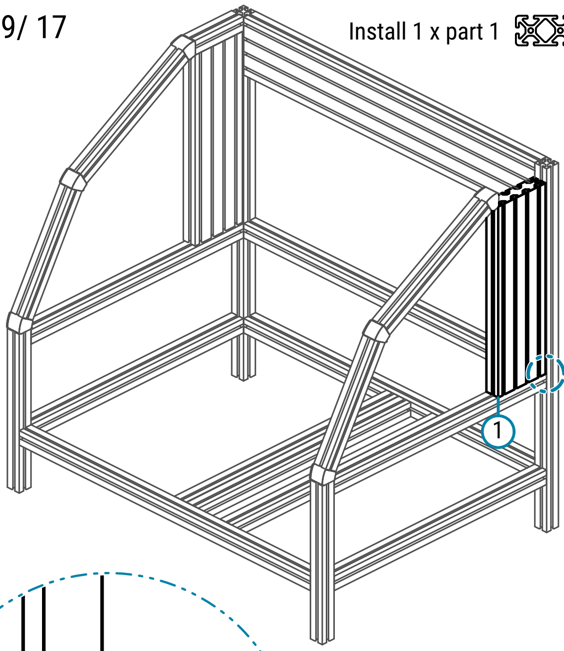

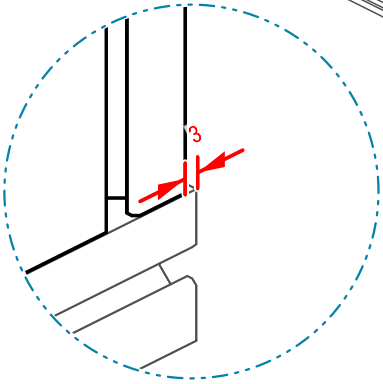

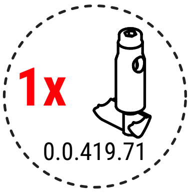

#### Step 10/ 17

Install 1 x part 6

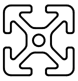

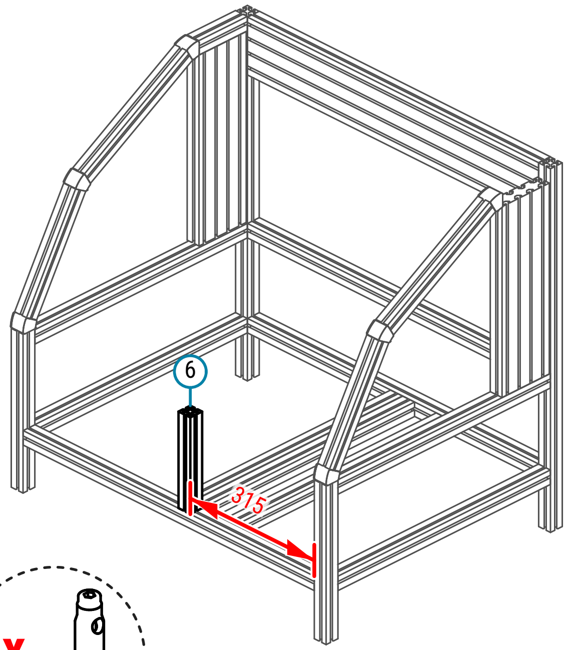

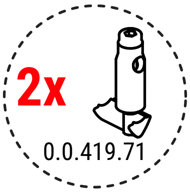

#### Step 11/ 17

Install 4 x part 3v

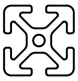

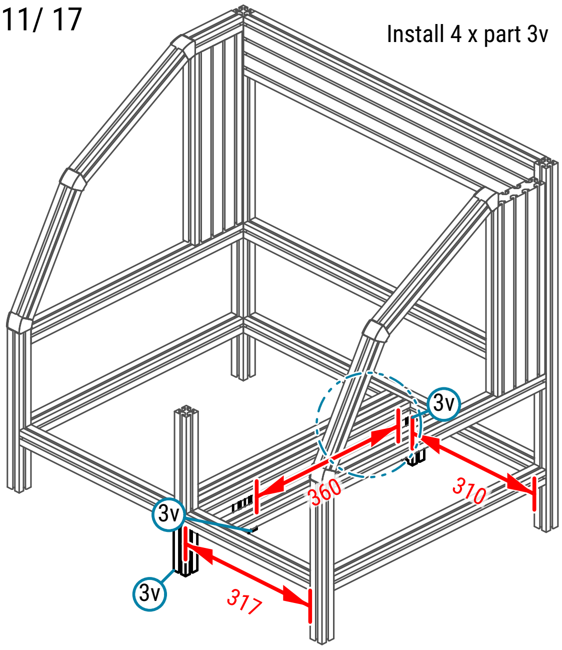

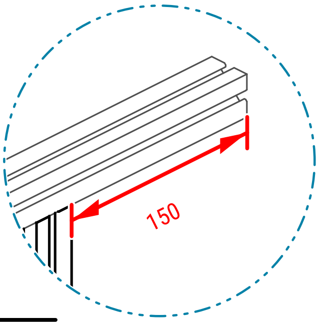

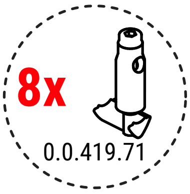

#### Step 12/ 17

Install 1 x part 5

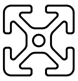

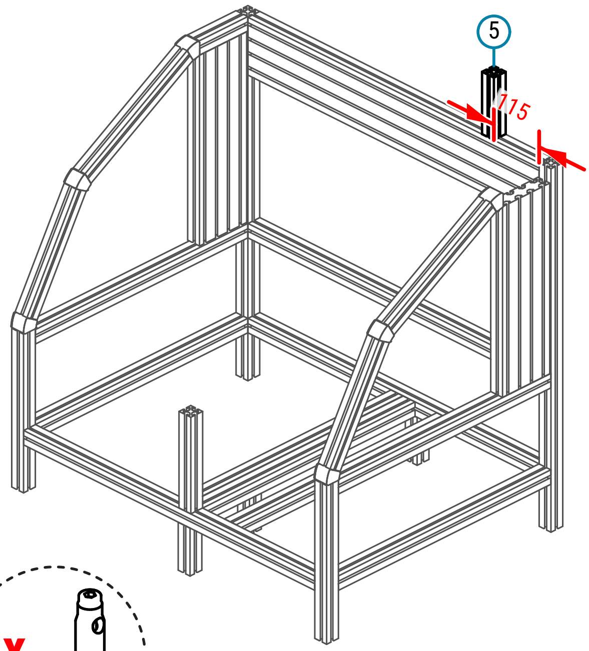

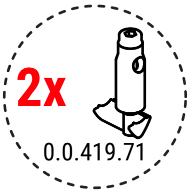

#### Step 13/ 17

Install 1 x part 9

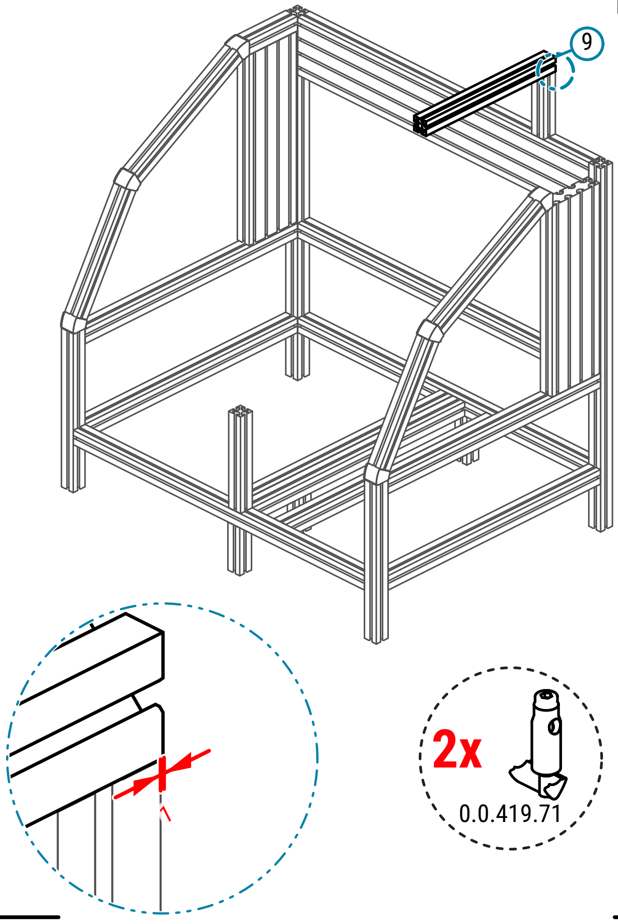

#### Step 14/ 17

Install 1 x part 4

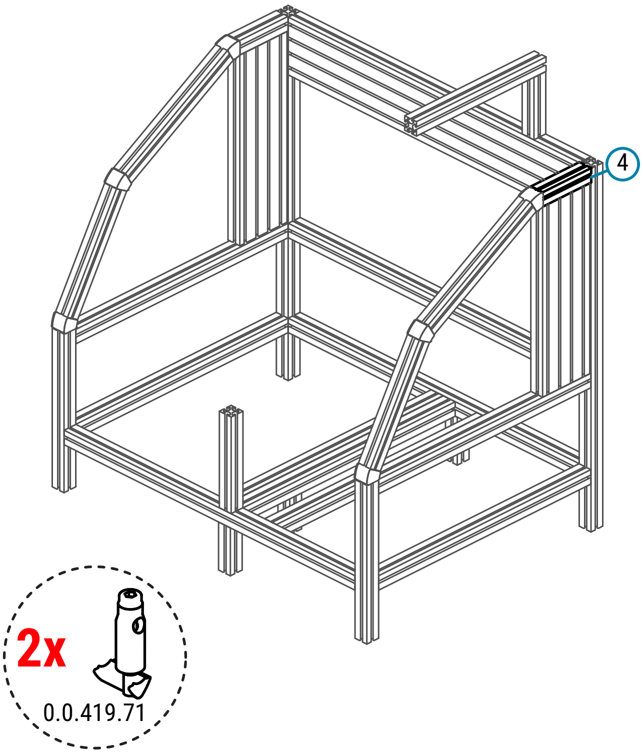

#### Step 15/ 17

Install 1 x part 11

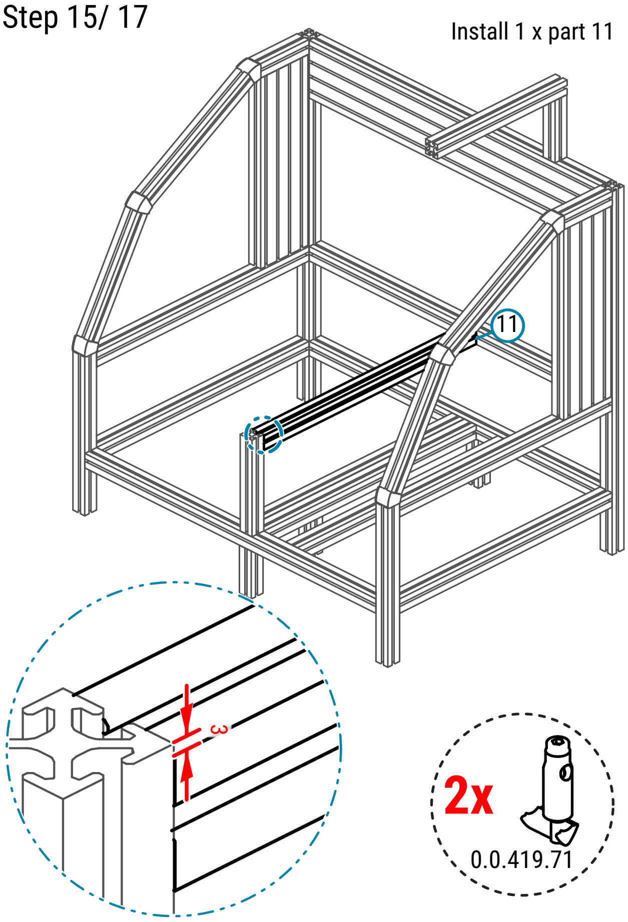

#### Step 16/ 17

Install 8 x part 17

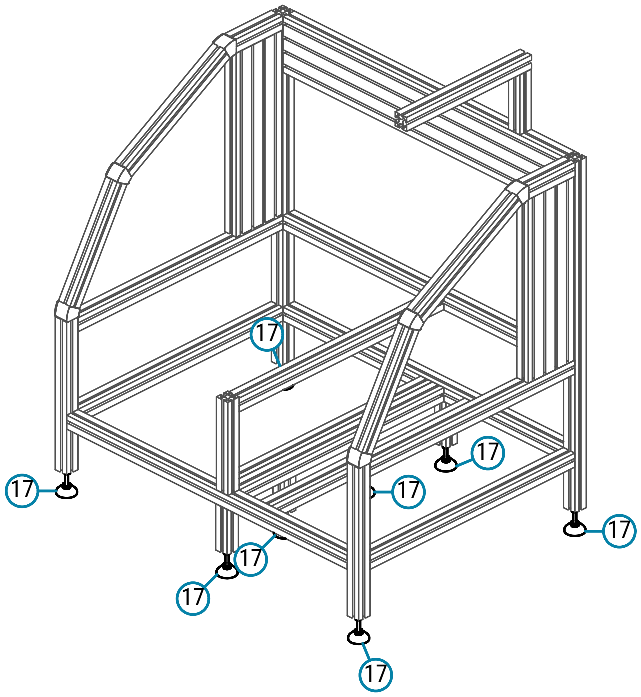

Step 17/ 17

Install 5 x part 15

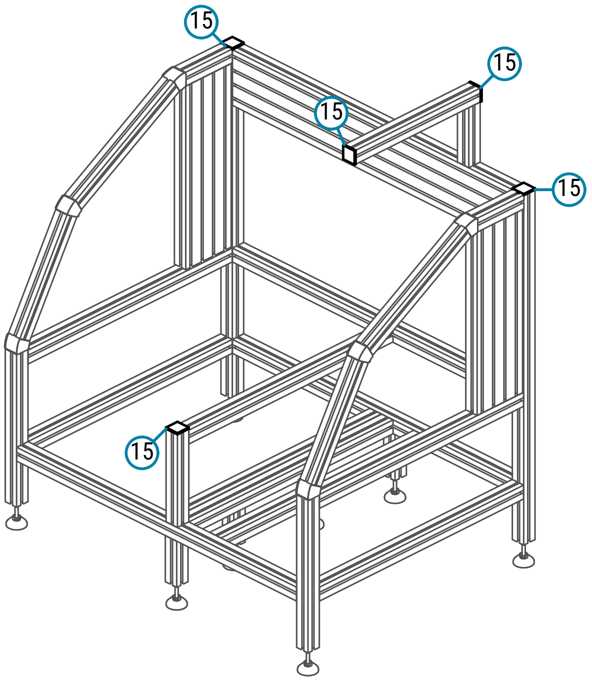
